# Crystallographic and thermodynamic evidence of negative cooperativity of flavin and tryptophan binding in the flavin-dependent halogenases AbeH and BorH

**DOI:** 10.1101/2023.08.22.554356

**Authors:** Md Ashaduzzaman, Kazi Lingkon, Aravinda J. De Silva, John J. Bellizzi

## Abstract

The flavin-dependent halogenase AbeH produces 5-chlorotryptophan in the biosynthetic pathway of the chlorinated bisindole alkaloid BE-54017. We report that *in vitro*, AbeH (assisted by the flavin reductase AbeF) can chlorinate and brominate tryptophan as well as other indole derivatives and substrates with phenyl and quinoline groups. We solved the X-ray crystal structures of AbeH alone and complexed with FAD, as well as crystal structures of the tryptophan-6-halogenase BorH alone, in complex with 6-chlorotryptophan, and in complex with FAD and tryptophan. Partitioning of FAD and tryptophan into different chains of BorH and failure to incorporate tryptophan into AbeH/FAD crystals suggested that flavin and tryptophan binding are negatively coupled in both proteins. ITC and fluorescence quenching experiments confirmed the ability of both AbeH and BorH to form binary complexes with FAD or tryptophan and the inability of tryptophan to bind to AbeH/FAD or BorH/FAD complexes. FAD could not bind to BorH/tryptophan complexes, but FAD appears to displace tryptophan from AbeH/tryptophan complexes in an endothermic entropically-driven process.

## Introduction

Flavin-dependent halogenases (FDHs; EC 1.14.19.9, 1.14.19.58, 1.14.19.59) evolved to install halogen atoms in the biosynthetic pathways of mostly bacterial and fungal natural products(1-4). They have attracted attention for their potential as biocatalytic tools for green organic synthesis due to their ability to regioselectively produce aryl halides that can be used for transition metal catalyzed cross coupling reactions(5-8). Challenges to synthetic applications of FDHs include limited substrate scope and suboptimal stability and catalytic efficiency, and efforts to overcome these hurdles have included mechanistic modeling using molecular dynamics simulation(9-11), protein engineering (12-14) and bioinformatic analysis and bioprospecting of diverse microbial genomes and metagenomes to discover new FDHs with different halide preferences, substrate tolerances, regioselectivities, and thermostabilities(15-20).

AbeH (UniProtKB F6LWA5) and BorH (UniProtKB M9QSI0) are FDHs encoded in soil actinomycete-derived gene clusters responsible for the biosynthesis of the chlorinated indolotryptoline natural products BE-54017(21) (*abe*) (GenBank JF439215) and borregomycin-A(22) (*bor*) (GenBank AGI62217), which were identified from environmental DNA libraries collected from Anza-Borrego desert soil (Figure 1). Based on sequence homology to FDHs and the location of chlorine atoms in BE-54017 and borregomycin A, which are assembled by oxidative dimerization of chlorinated *L*-tryptophan (Trp), AbeH was predicted to be a Trp-5- halogenase and BorH was predicted to be a Trp-6-halogenase. The gene clusters each contain an ORF predicted to encode a short-chain flavin reductase (FR), and it was hypothesized that AbeF (UniProtKB F6LWA7) and BorF (UniProtKB M9QXS1) are the FRs responsible for supplying AbeH and BorH, respectively with FADH_2_ (Figure 1).

**Figure 1:**
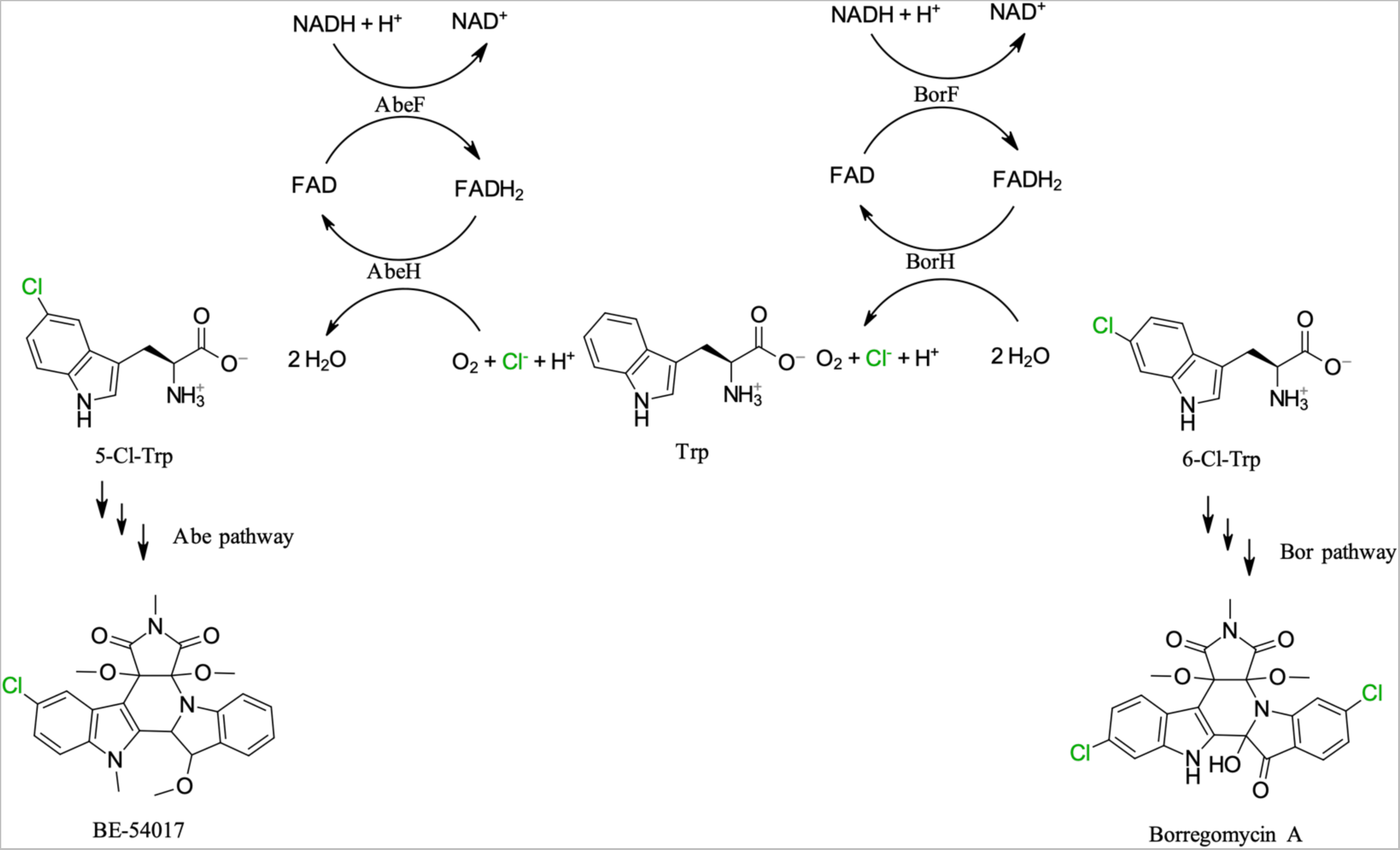
AbeH and BorH are flavin-dependent tryptophan halogenases. AbeH uses FADH_2_, O_2_ and Cl^-^ to convert Trp to 5-Cl-Trp and AbeF reduces the FAD released by AbeH to FADH_2_ using NADH to complete the catalytic cycle. Similarly, BorH uses FADH_2_ generated by BorF to convert Trp to 6-Cl-Trp. The other enzymes encoded by the *abe* and *bor* gene clusters convert 5-Cl-Trp and 6-Cl-Trp to the bisindole alkaloids BE-54017 and borregomycin A, respectively.

We have previously confirmed that BorF and AbeF are FRs that catalyze the NADH-dependent reduction of FAD to FADH_2_(23) and shown that BorH can brominate and chlorinate Trp at C6, and can also halogenate anthranilamide, 6-aminoquinoline, kynurenine, tryptamine, and 3-indolepropionic acid in the presence of BorF (24). We also solved the crystal structure of BorH/Trp (6UL2) to reveal Trp bound in the same orientation previously seen in the structure of the Trp-6-FDH Thal (6H44)(13), confirming that substrate orientation was the structural basis for its regioselectivity(24). In this work, we report the *in vitro* characterization of the Trp-5-halogenase activity of AbeH, the crystal structures of AbeH with and without bound FAD, and crystal structures of apo-BorH and BorH in complexes with FAD, Trp, and 6-Cl-Trp.

Despite extensive efforts, we were unsuccessful in obtaining crystal structures of ternary FDH/FAD/Trp complexes for either AbeH or BorH, though such complexes have been previously reported for RebH, PrnA, and PyrH (25-27). Our inability to solve the ternary complex structures, along with previous reports of negative cooperativity of flavin and Trp binding in RebH and Thal (13, 27, 28), motivated us to investigate the binding of FAD and Trp to AbeH and BorH using ITC and fluorescence quenching. Our results revealed an inability of AbeH and BorH to simultaneously bind FAD and Trp and suggest that the two halogenases have two distinct modes of negative cooperativity.

## Results

### AbeH is a tryptophan-5-halogenase

AbeH with an N-terminal hexahistidine tag was overexpressed in *E. coli* and purified to homogeneity by immobilized metal affinity, anion exchange, and size exclusion chromatography (Figure S1). AbeH chlorinated Trp in the presence of AbeF, FAD, NADH, and NaCl (Figure S2) as determined by RP-HPLC analysis of reaction mixtures monitoring A_280_ for disappearance of Trp (*t_R_* = 10.3 min) and appearance of a new peak (*t_R_* = 11.3 min) which has the same retention as a 5-Cl-Trp standard (Figure S2). ESI-MS analysis of this peak (*m/z*: 239.0574 and 241.0559) confirmed the product was monochlorinated Trp based on mass and isotopic ratio (Table S1), and _1_H NMR verified that the isomer produced by AbeH is 5-Cl-Trp, verifying the predicted regioselectivity based on the position of the Cl in BE-54017 (Figure S3). Substituting NaBr for NaCl resulted in suppression of the 5-Cl-Trp peak and the appearance of a later-eluting peak (*t_R_* = 11.5 min) matching the retention time of a 5-Br-Trp standard (Figure S2), and the identity of this product was confirmed by ESI-MS (*m/z:* 283.0103, 285.0078 in 1:1 ratio). No product peak was observed when NaCl was replaced with NaI. The specific activity of AbeH for chlorination of Trp is approximately 2.5x the specific activity for Trp bromination (87 and 34 µM min^-1^ mg^-1^, respectively) (Figure S4).

### AbeH and BorH chlorinate and brominate indole, benzene, and quinoline derivatives

We investigated the halogenation activity of AbeH against 20 aromatic substrates and found that AbeH could chlorinate and brominate compounds with different aromatic scaffolds (phenyl, indole, and quinoline) (Table 1, Table S1, and Figure S5). AbeH halogenated indole and 3-indolepropionic acid, but showed no halogenation activity against 5-cyanoindole, 5- hydroxytryptopan, and serotonin, presumably because the C5 position was already substituted. In addition to the indole derivatives, AbeH halogenated anthranilamide, but not anthranilic acid, aniline, benzamide, 3-aminobenzamide, or 4-aminobenzamide, suggesting both the amino group and amide are required they must be in a 1,2 relative orientation. Similarly, 6-aminoquinoline and 7-aminoquinoline were chlorinated and brominated by AbeH, but quinoline, 5-aminoquinoline, 6- aminoquinoline, and 8-aminoquinoline were not. Though AbeH prefers tryptophan chlorination over bromination, for all other halogenated products, the brominated product was produced in higher yield than the chlorinated product under the same reaction conditions (Table 1, Table S1, and Figure S5).

**Table 1:**
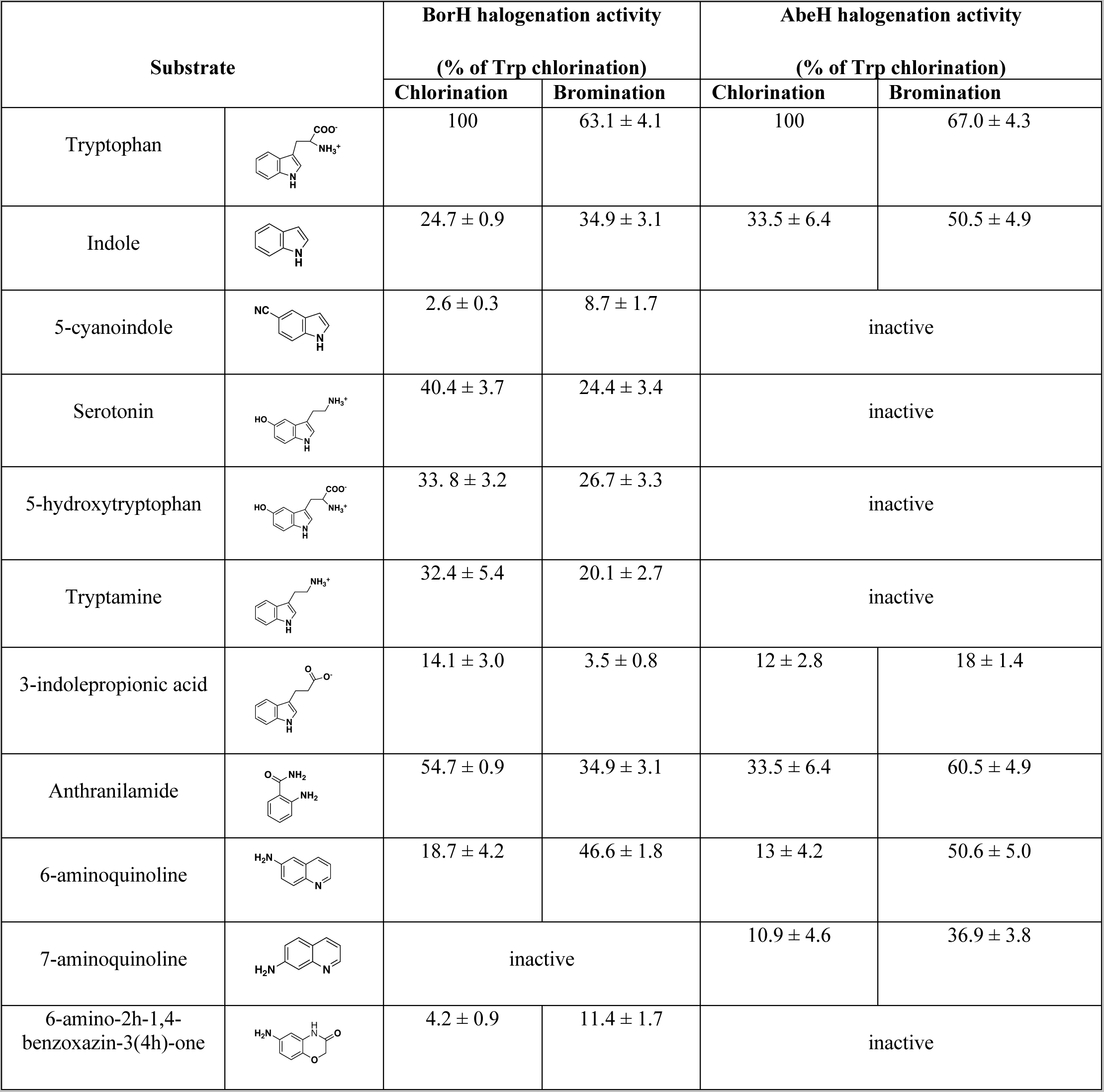
Relative activities for halogenation of aromatic substrates by BorH and AbeH.

| Substrate |  | BorH halogenation activity |  | AbeH halogenation activity |  |
| --- | --- | --- | --- | --- | --- |
|  |  | (% of Trp chlorination) |  | (% of Trp chlorination) |  |
|  |  | Chlorination | Bromination | Chlorination | Bromination |
| Tryptophan |  | 100 | 63.1 ± 4.1 | 100 | 67.0 ± 4.3 |
| Indole |  | 24.7 ± 0.9 | 34.9 ± 3.1 | 33.5 ± 6.4 | 50.5 ± 4.9 |
| 5-cyanoindole |  | 2.6 ± 0.3 | 8.7 ± 1.7 | inactive |  |
| Serotonin |  | 40.4 ± 3.7 | 24.4 ± 3.4 | inactive |  |
| 5-hydroxytryptophan |  | 33.8 ± 3.2 | 26.7 ± 3.3 | inactive |  |
| Tryptamine |  | 32.4 ± 5.4 | 20.1 ± 2.7 | inactive |  |
| 3-indolepropionic acid |  | 14.1 ± 3.0 | 3.5 ± 0.8 | 12 ± 2.8 | 18 ± 1.4 |
| Anthranilamide |  | 54.7 ± 0.9 | 34.9 ± 3.1 | 33.5 ± 6.4 | 60.5 ± 4.9 |
| 6-aminoquinoline |  | 18.7 ± 4.2 | 46.6 ± 1.8 | 13 ± 4.2 | 50.6 ± 5.0 |
| 7-aminoquinoline |  | inactive |  | 10.9 ± 4.6 | 36.9 ± 3.8 |
| 6-amino-2h-1,4-benzoxazin-3(4h)-one |  | 4.2 ± 0.9 | 11.4 ± 1.7 | inactive |  |

We tested BorH against the same set of substrates as AbeH (Table 1, Table S1, and Figure S6). As previously observed(24), BorH halogenated a wide range of indole derivatives as well as anthranilamide and 6-aminoquinoline, but unlike AbeH, it could not halogenate 7-aminoquinoline. BorH could halogenate 6-amino-2H-1,4-benzoxazin-3(4H)-one, which AbeH did not halogenate. BorH showed a higher rate of bromination than chlorination for indole, 5-cyanoindole, 6- aminoquinoline, and 6-amino-2H-1,4-benzoxazin-3(4H)-one, but a higher yield of chlorination than bromination for the other substrates tested.

### Crystal structures of AbeH/FAD/Cl^-^ and apo-AbeH

We grew diffraction-quality single crystals of AbeH in two crystal forms (Table 2). Orthorhombic AbeH crystals grew only in the presence of FAD and crystallized in space group *P*2_1_2_1_2_1_ with two AbeH/FAD/Cl^-^ complexes in the asymmetric unit (ASU). Apo-AbeH crystallized in space group *P*2_1_ with four molecules in the ASU.

**Table 2:** X-ray diffraction data and refinement statistics for AbeH crystal structures.

| Parameters | Apo-AbeH<br>(PDB:8FOX) | AbeH/FAD/Cl<sup>-</sup><br>(PDB:8FOV) |
| --- | --- | --- |
| **Wavelength (Å)** | 0.97857 | 0.97872 |
| **Beamline** | APS 21-ID-G | APS 21-ID-F |
| **Resolution range (Å)** | 60.1-1.89 (1.958-1.89) | 53.91-1.86 (1.926-1.86) |
| **Space group** | *P*2<sub>1</sub> | *P*2<sub>1</sub>2<sub>1</sub>2<sub>1</sub> |
| **Unit cell (Å, °)** | 63.4496, 251.13, 68.01<br>90, 108.71, 90 | 56.71, 112.29, 173.59<br>90, 90, 90 |
| **Total reflections** | 319567 (31371) | 186383 (18541) |
| **Unique reflections** | 159987 (15847) | 93637 (9276) |
| **Multiplicity** | 2.0 (2.0) | 2.0 (2.0) |
| **Completeness (%)** | 99.88 (99.37) | 99.63 (99.96) |
| **Mean $\langle I/\sigma(I) \rangle$** | 9.27 (2.88) | 7.10 (2.06) |
| **Wilson B-factor** | 18.52 | 23.36 |
| **R<sub>merge</sub>** | 0.04645 (0.274) | 0.06443 (0.3797) |
| **R<sub>meas</sub>** | 0.06569 (0.3875) | 0.09112 (0.537) |
| **R<sub>pim</sub>** | 0.04645 (0.274) | 0.06443 (0.3797) |
| **CC<sub>1/2</sub>** | 0.996 (0.8) | 0.99 (0.708) |
| **CC\*** | 0.999 (0.943) | 0.998 (0.911) |
| **Reflections used in refinement** | 159921(15845) | 93623 (9274) |
| **Reflections used for R<sub>free</sub>** | 2001 (208) | 894 (89) |
| **R<sub>work</sub>** | 0.1588(0.2149) | 0.1633(0.2414) |
| **R<sub>free</sub>** | 0.1834(0.2593) | 0.1960(0.2634) |
| **CC(work)** | 0.965 (0.896) | 0.966 (0.851) |
| **CC(free)** | 0.961 (0.853) | 0.938 (0.866) |
| **Number of non-hydrogen atoms** | 17945 | 8945 |
| **macromolecules** | 15940 | 7983 |
| **ligands** | 60 | 184 |
| **solvent** | 1945 | 778 |
| **Protein residues** | 1985 | 995 |
| **RMS(bonds)** | 0.003 | 0.012 |
| **RMS(angles)** | 0.65 | 1.18 |
| **Ramachandran favored (%)** | 97.14 | 96.24 |
| **Ramachandran allowed (%)** | 2.86 | 3.76 |
| **Ramachandran outliers (%)** | 0 | 0 |
| **Rotamer outliers (%)** | 0.24 | 0.6 |
| **Clash score** | 3.36 | 3.32 |
| **Average B-factor** | 22.6 | 28.94 |
| **macromolecules** | 21.37 | 28.13 |
| **ligands** | 36.09 | 31.66 |
| **solvent** | 32.22 | 36.58 |
| **Number of TLS groups** | 23 | 15 |

The AbeH/FAD/Cl^-^ (Figure 2A) structure was solved at 1.86 Å resolution using molecular replacement with Th-Hal (29)(5LV9.pdb, 66% sequence identity to AbeH as the search model. AbeH comprises 16 α-helices, 7 3_10_-helices, 1 π-helix and 24 β-strands, and as expected has the fold originally reported for PrnA(28, 30) and since seen in numerous other FDH structures(13, 24, 27, 30-32), consisting of a box-shaped subdomain with a Rossmann-like fold containing the flavin binding site with an appended pyramidal subdomain (Figure 2A). In other FDHs, the substrate binding site has been shown to lie at the interface between the box and the pyramid. Lys75 and Glu352 superimpose with the catalytic Lys and Glu residues identified in other FDHs structures. Both chains in the AbeH/FAD/Cl^-^ structure have a disordered region (Gly148-Gln160, between η3 and β9 in the box subdomain), which in other FDHs forms a lid over the Trp binding site when occupied (see discussion). Chain A has a second missing segment (Asp261-Arg265, between β16 and bβ17). This region forms α-helix α8 between β15 and β16 in chain B. There are also two residues missing from the C-terminus of both chains. The interface between the two chains involves 42 residues from each chain and buries approximately 1610 Å^2^. The two pyramidal subdomains nest together, aligning the rectangular box-shaped subdomains approximately parallel to one another. This dimer interface is the same as that observed in other FDHs structures.

**Figure 2:**
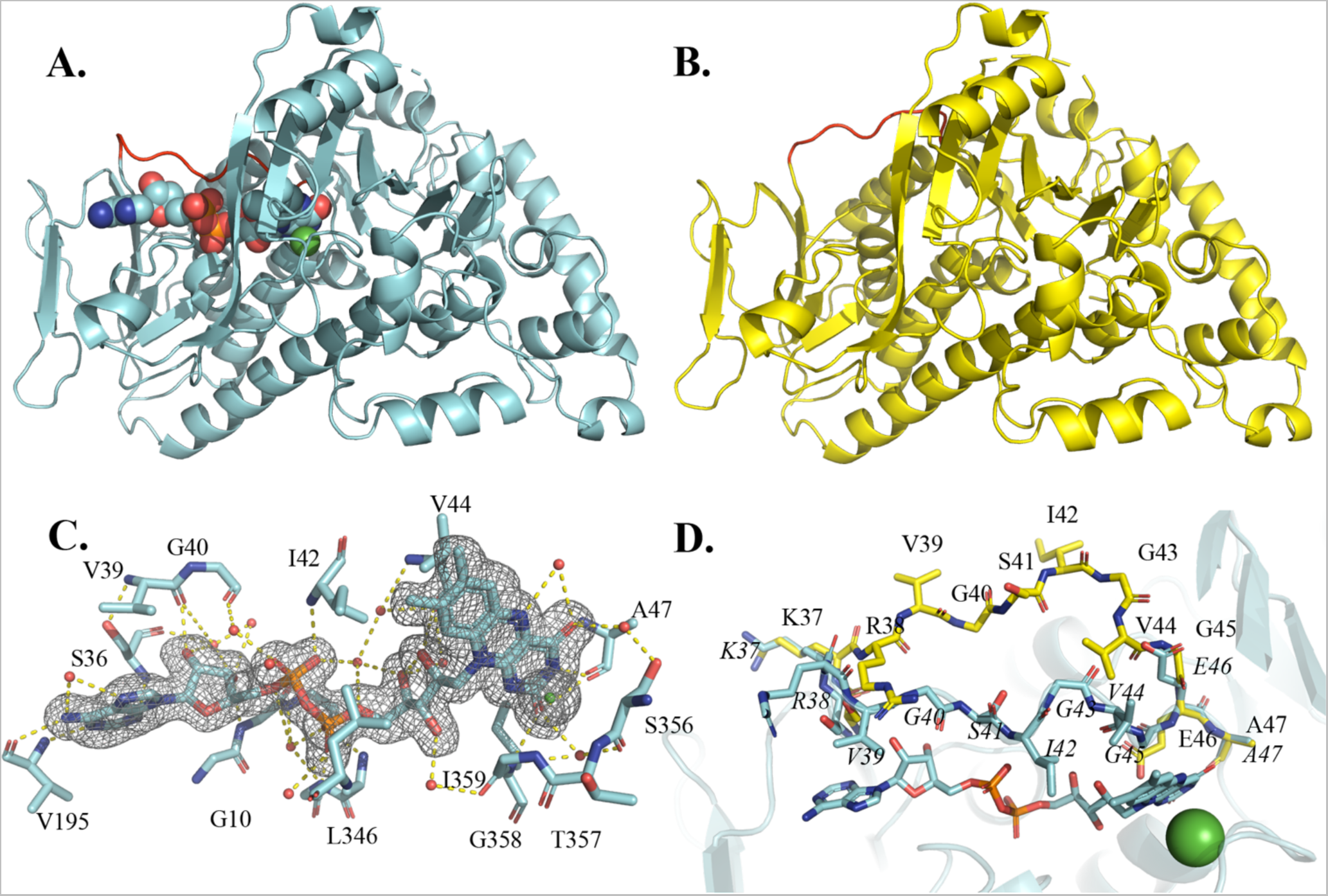
X-ray crystal structures of AbeH show two conformational states of the flavin binding loop. **A.** Ribbon diagram of chain A of AbeH/FAD/Cl^-^(cyan). FAD is shown as a space filling model, and Cl- is a green sphere. The flavin binding loop (in red) is in the closed conformation. **B.** Ribbon diagram of chain C of Apo-AbeH (yellow). The flavin binding loop (in red) is in the open conformation. **C.** Flavin binding site of chain A of AbeH/FAD/Cl^-^(cyan). FAD and residues contacting FAD are shown as sticks, Cl^-^ as a green sphere, water molecules as red spheres, and hydrogen bonds as dashed lines. The mesh is a Polder *F_o_-F_c_* omit map contoured at 3σ with FAD and Cl^-^ omitted from the map calculation. **D.** Superposition of the FAD binding sites of apo-AbeH chain C (yellow) and AbeH/FAD/Cl- chain A (cyan), illustrating the open and closed conformations of the flavin binding loop.

Density for FAD could be clearly seen and modeled in both chains (Figure 2C). A Cl^-^ ion was modeled in each chain in a strong spherical density peak adjacent to the *re*-face of the isoalloxazine ring of FAD; if modeled as water instead of Cl^-^, difference maps showed residual positive density in this peak. This site corresponds to the halide binding site identified in other FDHs structures (25). FAD binds to AbeH in the same manner seen in previously determined FDH/FAD structures. The isoalloxazine occupies a hydrophobic pocket with the *re*-face packed against η6 and the *si* face covered by the closed conformation of the flavin binding loop (located between β2 and α2) (Figure 2C, 2D). FAD makes hydrogen bonds with backbone atoms of Ala47, Ser 356, Ile359 (isoalloxazine), Ala13, Ile42, Leu346 (pyrophosphate), Gly10, Gly11, Ser36, and Val195 (adenosine) and 12 water-mediated hydrogen bonds (Figure 2C). The Cl^-^ is 3.2 Å above the *re*-face of the isoalloxazine, forming ion-dipole interactions with the backbone amide groups of Thr357 and Gly358 (Figure 2C).

The orthorhombic crystal used to solve the AbeH/FAD/Cl^-^ structure crystallized from a solution containing AbeH, FAD, and Trp, but there was no density observed in the expected Trp binding site. Data were collected on 12 more AbeH/FAD orthorhombic crystals that were either grown in the presence of Trp or soaked in Trp at concentrations from 2.5-40 mM for up to 3 days, but none of the maps generated contained density in the Trp binding site.

The overall Cα root mean square deviation (rmsd) between the two chains of AbeH/FAD/Cl^-^ is 0.47 Å. The major differences between the two chains are the lack of density for α8 and surrounding residues between β16 and β17 in chain A, and a difference in the orientation of the peptide bond between Val39 and Gly40 in the flavin binding loop. In chain A, the carbonyl of Val39 is pointed towards FAD and accepts a hydrogen bond from the ribose C2’-OH, whereas in chain B, the peptide bond is flipped and the amide of Gly40 donates a hydrogen bond to the ribose C2’-OH.

The apo-AbeH structure was solved from a 1.89 Å dataset collected from a monoclinic AbeH crystal using molecular replacement with chain B of the AbeH/FAD/Cl^-^ structure as a search model (Figure 2B). There are four AbeH chains in the asymmetric unit, arranged as two dimers (B/D and A/C) with the same dimer interface seen in the AbeH/FAD/Cl^-^ structure. The monoclinic crystal used to solve the apo-AbeH structure had been soaked in Trp, but there was no electron density in the Trp binding site. The most significant backbone difference between the chains in the apo-AbeH structure and the AbeH/FAD/Cl^-^ structure is in the flavin binding loop, which is closed over FAD in both chains of the complex structure but is disordered in Chains A and B and in the open conformation in chains C and D in the apo-AbeH structure (Figure 2D). In addition, the peptide bond between Gly10 and Gly11 is flipped, so that in apo-AbeH the Gly10 carbonyl O is hydrogen bonded to the NH of Gly14, and in AbeH/FAD/Cl^-^, the Gly11 NH is engaged in a water-mediated interaction with the FAD pyrophosphoryl group (a similar flip was previously observed between the open and closed conformations of Thal (13)). In the apo-AbeH structure, all four chains are missing the η3-β9 segment, chains A, C and D are missing the segment between ×15 and ×16, and chains C and D are missing five additional residues at the C-terminus. The four apo-AbeH chains have Cα rmsd values ranging from 0.33-0.39 Å with one another and from 0.74-1.08 Å with chain B of the AbeH/FAD/Cl^-^ structure.

### Crystal structures of apo-BorH and BorH substrate and product complexes

We determined X-ray structures of BorH in the absence of substrates/products (apo-BorH), BorH complexed with product 6-Cl-Trp (BorH/6-Cl-Trp), and a structure containing two BorH/FAD complexes and two BorH/Trp complexes in the asymmetric unit (BorH/Trp + BorH/FAD) (Figure 3 and Table 3). All structures were crystallized as apo-BorH in the same crystal form we used to solve the BorH/Trp complex (space group *P*2_1_ with four BorH molecules in the ASU, packed as two dimers A/D and B/C with the same dimer interface seen in the AbeH structures)(24), and soaked with substrates/products before cryoprotection, freezing, and data collection. The BorH/Trp + BorH/FAD structure was solved from a crystal soaked in both FAD and Trp in an attempt to form a ternary complex.

**Figure 3:**
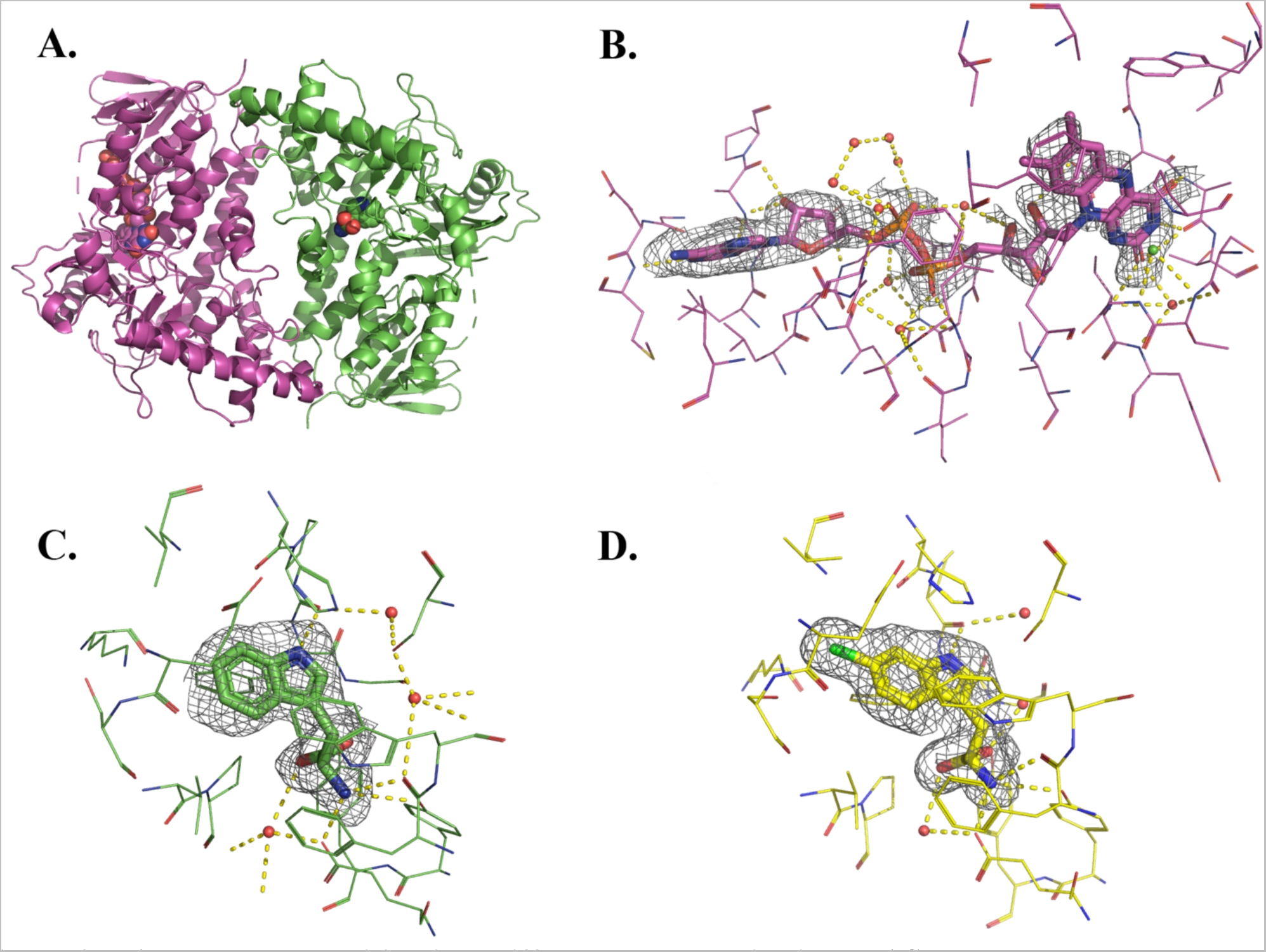
FAD and Trp partition into different BorH chains in the ASU. A. Ribbon diagram of crystallographic dimer of BorH/Trp (Chain A; green) with BorH/FAD/Cl^-^(Chain C; magenta) from BorH crystals soaked in both FAD and Trp. B. Flavin binding site of chain C of BorH/FAD/Cl^-^(magenta) from the BorH/FAD/Trp crystal structure. FAD and residues contacting FAD are shown as sticks, Cl-as a green sphere, water molecules as red spheres, and hydrogen bonds as dashed lines. The mesh is a Polder *F_o_-F_c_* omit map contoured at 3α with FAD and Cl-omitted from the map calculation. C. Trp binding site of chain A of BorH/Trp (green) from the BorH/FAD/Trp crystal structure. Trp and residues contacting Trp are shown as sticks, water molecules as red spheres, and hydrogen bonds as dashed lines. The mesh is a Polder *F_o_-F_c_* omit map contoured at 3α with the Trp substrate omitted from the map calculation. D. 6-Cl-Trp bound in the Trp binding site of Chain A of the BorH/6-Cl-Trp crystal structure (yellow). 6-Cl-Trp and residues contacting Trp are shown as sticks, water molecules as red spheres, and hydrogen bonds as dashed lines. The mesh is a Polder *F_o_-F_c_* omit map contoured at 3α with the 6-Cl-Trp product omitted from the map calculation.

**Table 3:** X-ray diffraction data and refinement statistics for BorH crystal structures.

| Parameters | BorH/Trp + BorH/FAD<br>(PDB:8TTI ) | BorH/6-Cl-Trp<br>(PDB: 8TTJ) | Apo-BorH<br>(PDB:8TTK ) |
| --- | --- | --- | --- |
| **Wavelength (Å)** | 1.033 | 1.033 | 1.033 |
| **Beamline** | APS 21-ID-D | APS 21-ID-D | APS 21-ID-D |
| **Resolution range (Å)** | 54.72 -1.98 (2.05 -1.98) | 34.98-1.98 (2.05 - 1.98) | 37.92-1.98 (2.05 - 1.98) |
| **Space group** | *P*2<sub>1</sub> | *P*2<sub>1</sub> | *P*2<sub>1</sub> |
| **Unit cell (Å, °)** | 73.68, 157.41, 112.83<br>90, 104.06, 90 | 73.52, 157.971, 112.58<br>90, 104.06, 90 | 73.74, 157.58, 112.97<br>90, 104.25, 90 |
| **Total reflections** | 312471 (31632) | 331395 (33446) | 321503 (32446) |
| **Unique reflections** | 163198 (16589) | 169702 (16993) | 164806 (16702) |
| **Multiplicity** | 1.9 (1.9) | 2.0 (2.0) | 2.0 (1.9) |
| **Completeness (%)** | 93.91 (93.15) | 98.24 (98.67) | 95.03 (96.50) |
| **Mean $\langle I/\sigma(I) \rangle$** | 3.74 (0.62) | 4.91 (1.06) | 5.59 (0.98) |
| **Wilson B-factor** | 30.89 | 27.45 | 26.22 |
| **R<sub>merge</sub>** | 0.1345 (1.316) | 0.0997 (0.7698) | 0.09669 (0.753) |
| **R<sub>meas</sub>** | 0.1902 (1.861) | 0.1401 (1.089) | 0.1367 (1.065) |
| **R<sub>pim</sub>** | 0.1345 (1.316) | 0.09909 (0.7698) | 0.09669 (0.753) |
| **CC<sub>1/2</sub>** | 0.98 (0.1) | 0.987 (0.247) | 0.957 (0.377) |
| **CC\*** | 0.995 (0.426) | 0.997 (0.629) | 0.989 (0.74) |
| **Reflections used in refinement** | 162282 (16029) | 169634 (16965) | 164625 (16680) |
| **Reflections used for R<sub>free</sub>** | 1989 (198) | 1996 (196) | 1993 (195) |
| **R<sub>work</sub>** | 0.2288 (0.3778) | 0.1982 (0.3326) | 0.2210 (0.3613) |
| **R<sub>free</sub>** | 0.2701 (0.4186) | 0.2362 (0.3864) | 0.2489 (0.4160) |
| **CC(work)** | 0.931 (0.457) | 0.950 (0.613) | 0.934 (0.678) |
| **CC(free)** | 0.913 (0.328) | 0.937 (0.604) | 0.923 (0.617) |
| **Number of non-hydrogen atoms** | 18159 | 18166 | 17760 |
| **macromolecules** | 16745 | 16784 | 16652 |
| **ligands** | 171 | 140 | 80 |
| **solvent** | 1243 | 1242 | 1028 |
| **Protein residues** | 2083 | 2084 | 2076 |
| **RMS(bonds)** | 0.003 | 0.005 | 0.003 |
| **RMS(angles)** | 0.50 | 0.72 | 0.61 |
| **Ramachandran favored (%)** | 97.04 | 97.43 | 96.93 |
| **Ramachandran allowed (%)** | 2.96 | 2.57 | 3.07 |
| **Ramachandran outliers (%)** | 0.00 | 0.00 | 0.00 |
| **Rotamer outliers (%)** | 0.70 | 0.29 | 1.29 |
| **Clash score** | 5.90 | 4.02 | 7.06 |
| **Average B-factor** | 44.20 | 37.97 | 39.65 |
| **macromolecules** | 44.15 | 37.74 | 39.62 |
| **ligands** | 51.27 | 56.75 | 52.90 |
| **solvent** | 43.97 | 38.95 | 39.09 |
| **Number of TLS groups** | 25 | 25 | 20 |

**Table 4.**
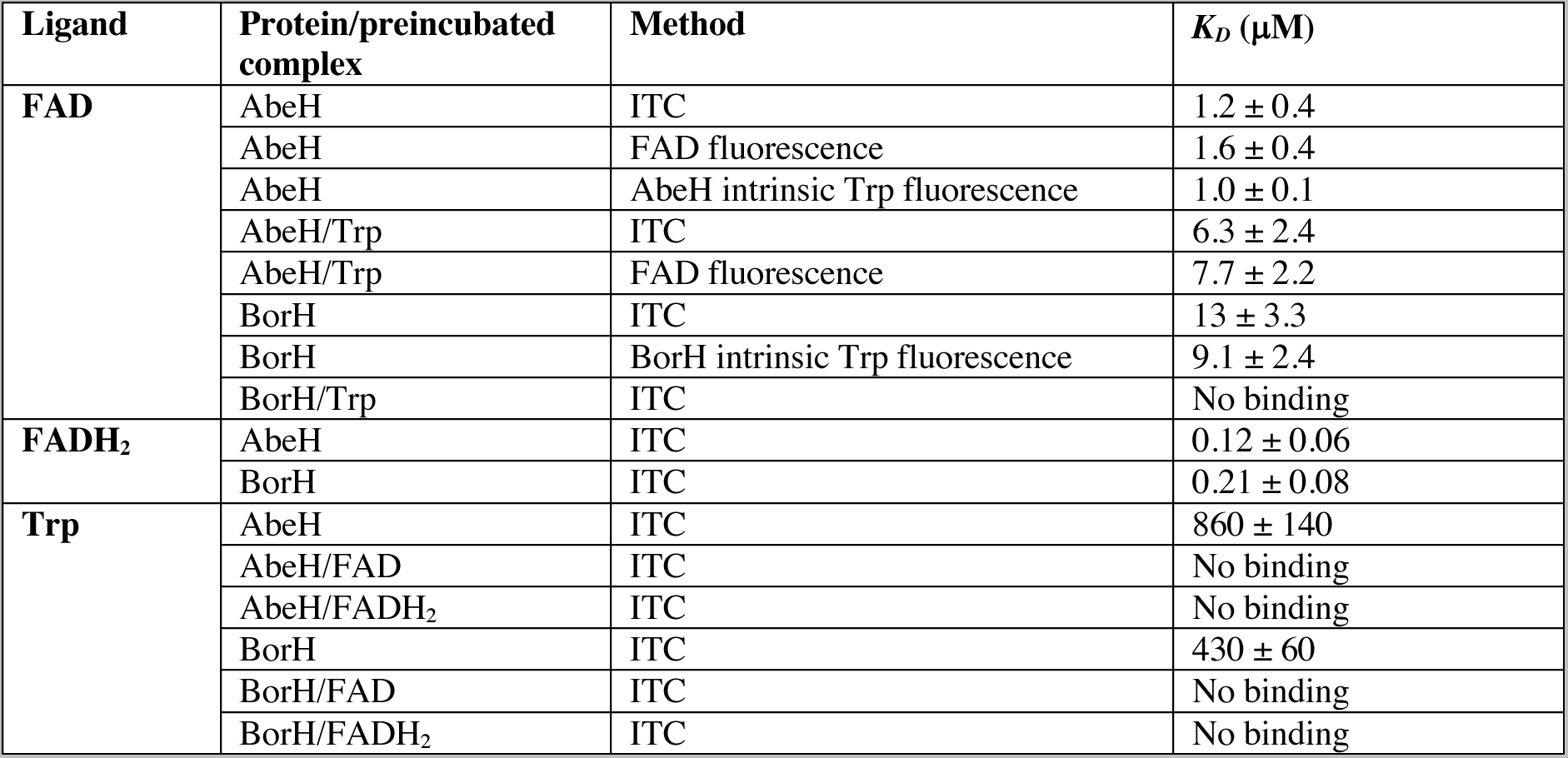
*K_D_* values determined for FAD, FADH_2_, and Trp binding to AbeH and BorH. Other ITC determined binding parameters are in **Table S2**.

There are no significant backbone differences among the 16 chains of the four BorH crystal structures. In our previous BorH/Trp structure (6UL2), there was no density for the flavin binding loop in chains B-D and weak but interpretable density that allowed us to model it in the open conformation in chain A. None of the chains in the apo-BorH, BorH/6-Cl-Trp, or BorH/Trp + BorH/FAD structures had density that enabled us to model the flavin binding loop, even in the chains where FAD is bound.

In the structure solved from the crystal soaked in both FAD and Trp, Chains A and B have Trp bound (but no FAD) and chains C and D have FAD bound (but no Trp) (Figure 3), arranged in the ASU as two heterodimers AD and BC (each containing BorH/Trp and BorH/FAD). The FAD bound to chains C and D has strong electron density for the adenosine and phosphate groups, with progressively weaker density for the ribityl and the isoalloxazine moiety. This is likely due to increased conformational mobility of the isoalloxazine and ribityl portions of FAD because the flavin binding loop is not in the closed conformation, and mirrors previous observations for Thal (33) in which the AMP was modeled but no other part of FAD was seen. FAD makes hydrogen bonds with backbone atoms of Ala50, Ile361 (isoalloxazine), Ala13, Ile42, Leu348 (pyrophosphate), Gly13, Thr15, Ala16, Gly17, and Met197 (adenosine) and 12 water-mediated hydrogen bonds. Spherical density at the putative halide binding site (adjacent to Thr359 and Gly360) was modeled as water, and unlike the AbeH/FAD/Cl^-^ structure, there was no residual positive density in difference maps.

The BorH/Trp interactions are unchanged when comparing the Trp bound chains (A and B) from the BorH/Trp + BorH/FAD structure to our previously-determined BorH/Trp structure (6UL2). The BorH/6-Cl-Trp structure (Figure 3D) has clear density for the product in all four chains and 6-Cl-Trp makes the same interactions with BorH as those previously observed in the BorH/Trp complex. The Cl^-^ of the 6-Cl-Trp is approximately 3 Å from Nε of the catalytic Lys79.

In chains without Trp or 6-Cl-Trp bound (BorH/FAD + BorH/Trp chains C, D, apo-BorH chains A-D), the density is weaker for the substrate binding lid (α14-α15, Val447-Glu460), suggesting that region becomes more highly ordered in the presence of substrate.

### Binding of Trp, FAD, and FADH_2_ to BorH and AbeH

We used isothermal titration calorimetry to analyze the binding of Trp, FAD, and FADH_2_ to AbeH and BorH (Figures 4-6 and S7, S8; Tables 4 and S2). The apparent *K_D_* for the AbeH/FAD complex was determined by ITC to be 1.2 µM (Figure 4A), which was consistent with values independently determined from fluorescence quenching titration experiments using FAD fluorescence (*K_D_* = 1.6 µM) (S10A) and AbeH intrinsic Trp fluorescence (*K_D_* = 1 µM) (Figure S9B). FAD binds to BorH with approximately 10-fold lower affinity than it binds to AbeH, with *K_D_* for the BorH/FAD complex = 13 µM by ITC (Figure 4D) and 10 µM by quenching of BorH intrinsic Trp fluorescence upon FAD titration (Figure S9A). FADH_2_ binds with higher affinity than FAD to both AbeH (Figure 4B) and BorH (Figure 4E), with *K_D_* = 0.1 µM for AbeH/FADH_2_ and *K_D_* = 0.2 µM for BorH/FADH_2_ as determined by ITC.

**Figure 4:**
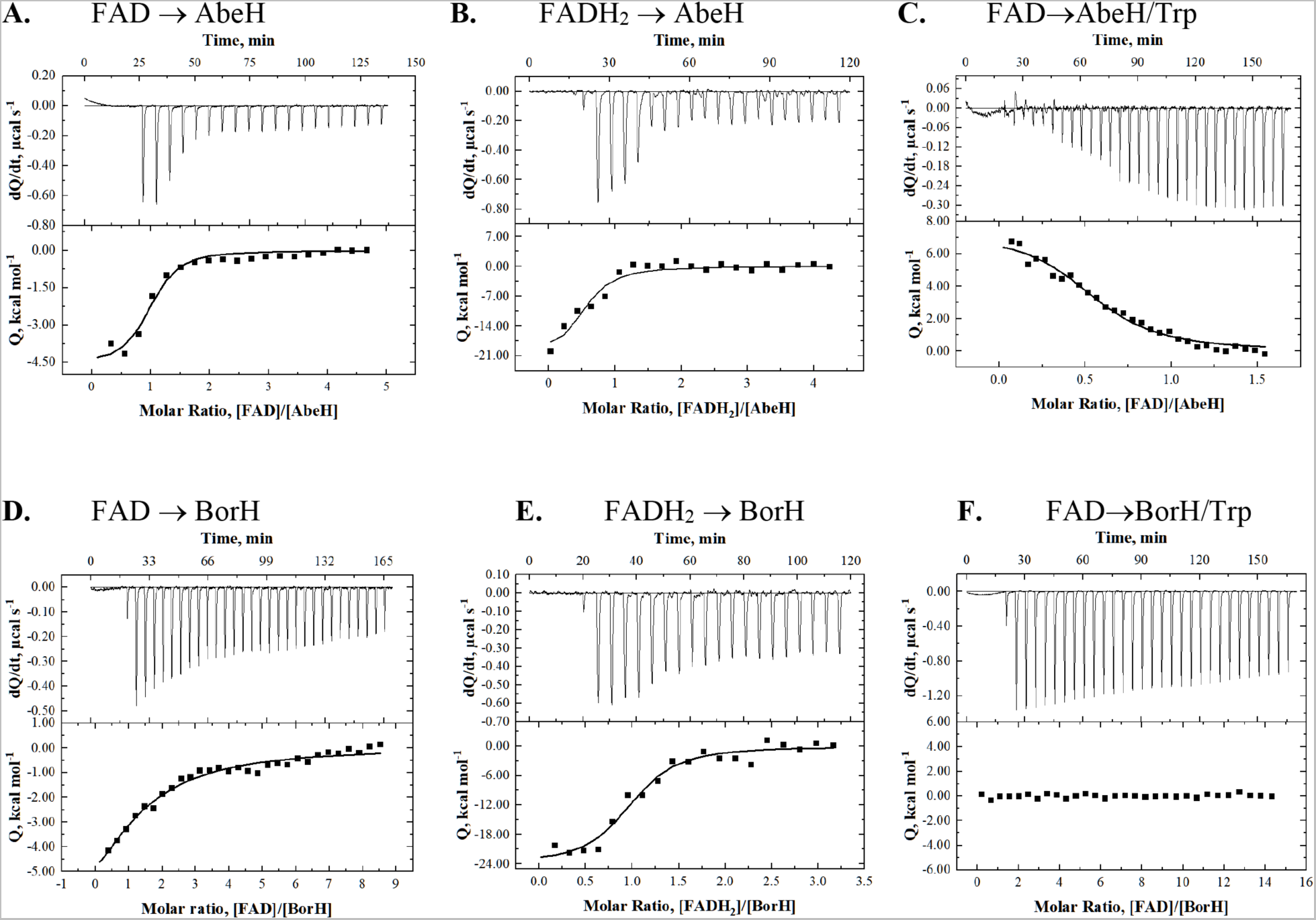
ITC analysis of FAD and FADH_2_ binding to AbeH and BorH. Top graph in each panel is the raw differential thermogram of titration with baseline as a solid line, and the bottom graph is the integrated titration curve showing corrected heat for each injection (^▄^) and best fit line ( ). **A**: Titration of FAD into AbeH. **B.** Titration of FADH_2_ into AbeH. **C**: Titration of FAD into AbeH/Trp. Binding was endothermic. **D:** Titration of FAD into BorH. **E** Titration of FADH_2_ into BorH. **F**: Titration of FAD into BorH/Trp (95% saturated). No binding was observed.

When Trp was titrated into either AbeH or BorH, heat of binding was only evolved only at high molar ratios of Trp to protein (Figure 5A, 5D, S7). Non-linear regression of ITC data for Trp binding to a one-site model in Origin was fixed to 1 to approximate the values for *K_A_*, *ΔH*, and *ΔS*. The calculated Trp affinities of AbeH and BorH are 500x and 30x lower than the FAD affinities (0.8 mM for Trp/AbeH and 0.4 mM for Trp/BorH).

**Figure 5:**
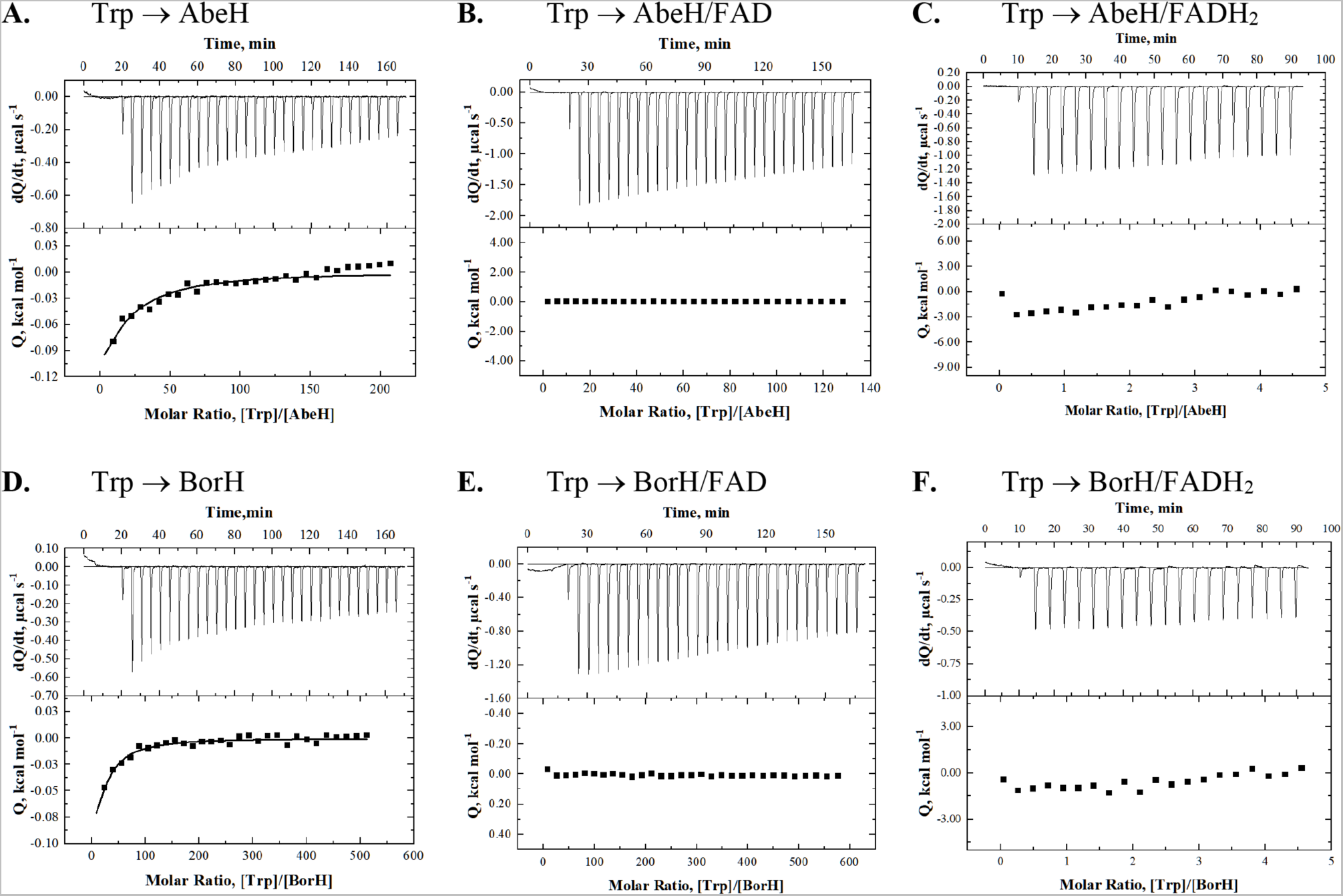
ITC analysis of Trp binding to AbeH and BorH. Top graph in each panel is the raw differential thermogram of titration with baseline as a solid line, and the bottom graph is the integrated titration curve showing corrected heat for each injection (^▄^) and best fit line ( ). **A**: Titration of Trp into AbeH. Binding was only observed at high molar ratio of Trp:AbeH **B.** Titration of Trp into AbeH/FADH_2_. No binding was observed. **C**: Titration of Trp into AbeH/FAD. No binding was observed. **D**: Titration of Trp into BorH. Binding was only observed at high molar ratio of Trp:BorH **E.** Titration of Trp into BorH/FADH_2_. No binding was observed. **F**: Titration of Trp into BorH/FAD. No binding was observed.

**Figure 6:**
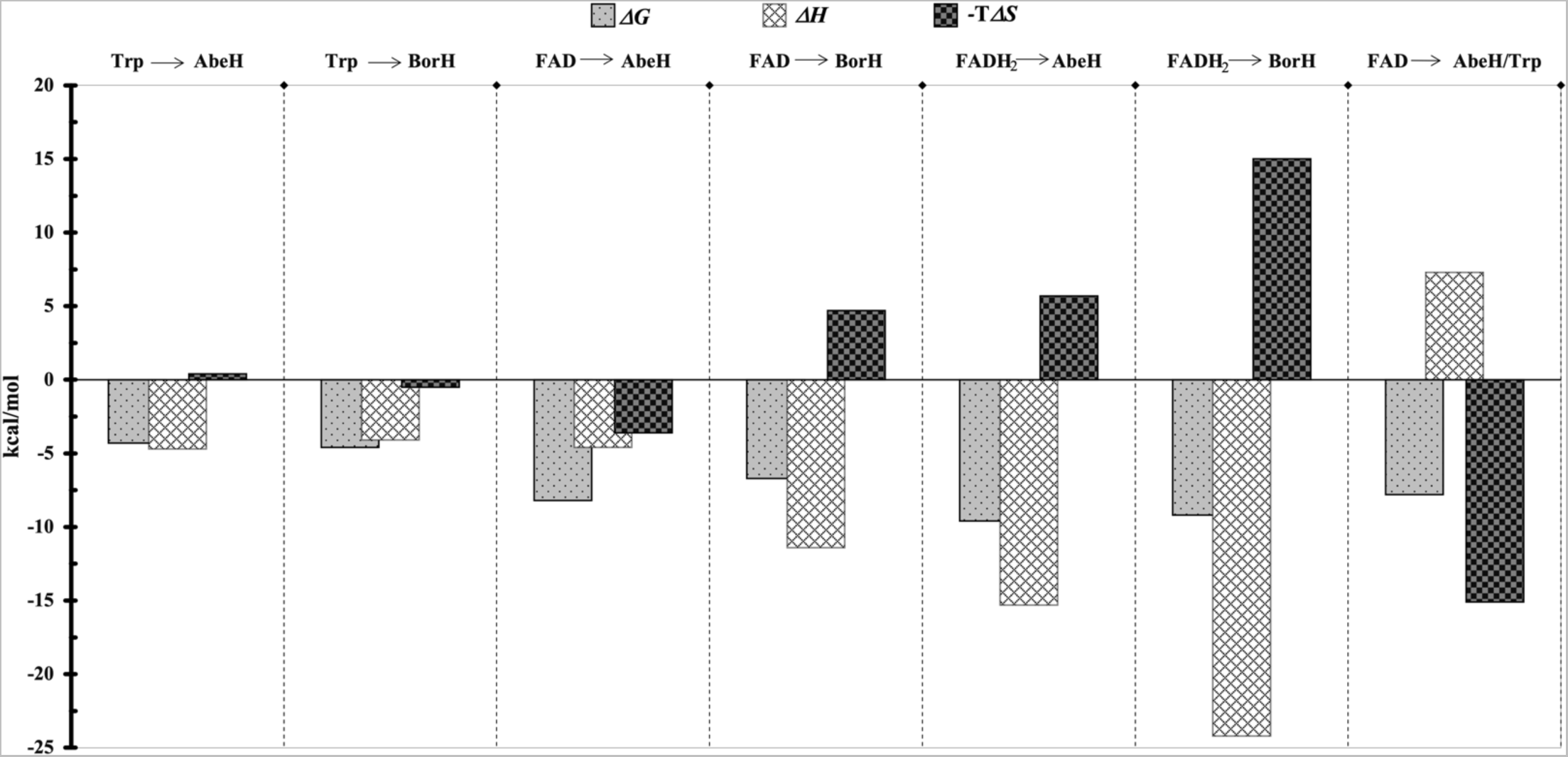
ITC-derived thermodynamic parameters. Values were obtained from non-linear least squares fit analysis of the ITC data using the one-site model in Origin 7.0. Other thermodynamic parameters are shown in Table S2.

Since we were unable to capture crystal structures of ternary FDH/FAD/Trp complexes, we tested whether we could observe the formation of ternary complexes by ITC (Figure S16). When AbeH or BorH was preincubated with FAD followed by titration with Trp, there was no evidence of Trp binding by ITC, even at very high molar ratios of Trp to protein (Figures 5B, 5E), indicating that AbeH/FAD and BorH/FAD complexes are unable to bind Trp. Likewise, titration of Trp into preincubated AbeH/FADH_2_ and BorH/FADH_2_ complexes showed no evidence of binding (Figure 5 C, 5F).

Titrating FAD into BorH preincubated with Trp also failed to detect any heat of binding, suggesting that BorH/Trp complexes are unable to bind FAD (Figure 4F), mirroring the partitioning observed in the crystal structure. In contrast, titrating FAD into AbeH preincubated with Trp resulted in heat being absorbed rather than released (Figure 4C). The apparent *K_D_* for FAD binding to AbeH in the presence of Trp determined by ITC is 2-5x higher than in the absence of Trp and is dependent on Trp concentration (*K_D_* = 2-6 µM, depending on the Trp: AbeH molar ratio). This dependence of apparent FAD affinity on Trp concentration was cross validated by fluorescence quenching titration (Figure S8 and S10).

## Discussion

Neither the orthorhombic AbeH crystals that grew in the presence of FAD nor the monoclinic crystals of AbeH that grew in the absence of FAD showed any evidence of substrate/product binding, despite dozens of attempts to incorporate Trp and 5-Cl-Trp by cocrystallization and soaking. Though Trp or 6-Cl-Trp could be soaked into the substrate binding site of monoclinic apo-BorH crystals, crystals soaked in both FAD and Trp resulted in a structure in which two molecules in the asymmetric unit had Trp bound, and two had FAD bound. Though crystal structures of ternary FDH/flavin/Trp complexes have been reported for RebH (2OA1.pdb, Chain B), PrnA (2AQJ.pdb, Chain A), and PyrH (2WET.pdb, Chain B) (25, 27), partitioning similar to our results has been seen previously for PyrH(30), RebH(27), and Thal(33) crystals, and FAD soaking difficulties have been reported for Xcc4156 apo-crystals(32). A model of negative coupling of FAD and Trp binding based on crystal structure analysis has been previously proposed for Thal(33). To investigate whether a similar phenomenon was occurring with AbeH and BorH, we carried out a detailed thermodynamic analysis of FDH cofactor and substrate binding. The ITC experiments were designed to test simultaneous binding of ligands (FAD or FADH_2_ and Trp) on ∼10 Å distant site of AbeH and BorH using a cooperativity model (Figure S16) described in the literature (34, 35).

Our ITC results are consistent with negative coupling, as we observed the formation of binary complexes between BorH or AbeH with FAD, FADH_2_ (with µM affinity) or Trp (with mM affinity) but saw no evidence of binding of Trp to AbeH/FAD or BorH/FAD or binding of FAD to BorH/Trp. Unexpectedly, titration of FAD into AbeH/Trp resulted in an endothermic binding event and the apparent *K_D_* for FAD binding to AbeH was slightly higher in the presence of Trp and increased with increasing Trp concentration. Fluorescence quenching titration confirmed that FAD can bind to AbeH but not BorH in the presence of saturating levels of Trp.

The binding of FAD to AbeH is enthalpically driven, but the binding of FAD to AbeH/Trp is entropically driven (Table S2). One interpretation of this result (consistent with the lack of any evidence of binding of Trp to AbeH/FAD) is that rather than a ternary complex being formed, FAD displaces Trp from the AbeH/Trp complex upon titration. The “lid” of the Trp binding site in AbeH and other members of its clade (Figure S11) is disordered in the absence of bound Trp, so ejection of Trp from the binding site upon FAD binding could increase the conformational entropy of AbeH. Conformational changes of the flavin-binding loop associated with rearrangements of water molecule networks have been described as a possible mechanism for formation of a Thal/FADH^-^ dead-end complex(11); water restructuring upon a shift from AbeH/Trp to AbeH/FAD could also plausibly account for some of the differences in *ΔH* and *ΔS* compared to titration of FAD into apo-AbeH.

The binding experiments were consistent with the inability of Trp and FAD to bind simultaneously to either BorH or AbeH, and suggested that FAD can displace Trp from AbeH, but not from BorH. This difference between the two proteins could be due to the differences in observed affinity for FAD and Trp of BorH and AbeH; BorH has lower affinity for FAD and higher affinity for Trp than does AbeH, meaning that the thermodynamic driving force for Trp displacement by FAD would be higher in AbeH than BorH. It could also mean that the structural basis for the negative cooperativity is different between BorH and AbeH. It is notable that AbeH and BorH have strikingly different architecture of the Trp binding site. Phylogenetic analysis of 21 different FDHs with solved crystal structures places AbeH and BorH in two different clades (Figure S11) with the AbeH clade including the Trp-5-halogenases AbeH and PyrH and the Trp-6-halogenases Th-Hal, SttH, and Tar14, and the BorH clade including the Trp-6-halogenases BorH and Thal and the Trp-7-halogenases PrnA and RebH(13, 24, 25, 27, 29–31, 36).

The AbeH clade and the BorH clade have marked differences in the architecture of the pyramidal subdomain resulting from insertions/deletions that alter the Trp binding site(24, 26)(Figures S12, S13). The lid covering the Trp binding site in AbeH/PyrH is part of the segment connecting α6 and β9 (residues 152-167). In the BorH/Thal clade, the connection between these two secondary structural elements is shorter, has a completely different topology, and does not form part of the Trp binding site. Instead, in the BorH/Thal clade, the lid comes from an insertion in the region between α14-α17 in BorH (Val447-Glu460), which contains two short perpendicular α-helices and is absent in the AbeH/PyrH clade (Figure S13). In both clades, the lid is less ordered in the absence of Trp, but the different topology of the two subfamilies means that the lid conformation may be coupled to different structural features.

Previous models for FDH negative coupling focus on the most obvious conformational change, the opening and closing of the flavin binding loop, and the effects that it has on residues immediately following the loop(30, 33). The flavin binding loop (Ser36-Glu46 in AbeH, Ala39- Glu49 in BorH) is usually (but not always) found in a closed conformation in structures with FAD or FADH_2_ bound and in an open conformation (or absent due to weak or missing electron density) in the absence of flavin. At the C-terminus of the flavin binding loop is a Glu (Glu46 in AbeH, Glu49 in BorH), the Cα of which acts as a pivot point about which the Glu side chain and peptide backbone exchange positions when the flavin binding loop moves from open (Glu side chain points towards vacant isoalloxazine binding site) to closed (Glu side chain flips out of binding site to make room for isoalloxazine).

Residues immediately C-terminal to the flavin binding loop (Phe 49/Ser50 in AbeH/PyrH, Val52/Pro53 in BorH/Thal) have been implicated in the inhibition of Trp binding, as they move into the Trp binding site when the flavin binding loop closes and move out when the loop opens, due to backbone shifts caused by the flipping of the adjacent Glu46/Glu49. This region is variously described as the Trp gate(37) and the Trp 1 region(38). Closing of the Trp gate can be seen when comparing the crystal structures of AbeH and PyrH. PyrH is the closest AbeH homolog (62% identity) that has been solved in a complex with Trp, and as it is also a Trp-5-FDH like AbeH and all of the residues contacting Trp in PyrH(30) are conserved in AbeH, it is a reasonable proxy for the elusive AbeH/Trp structure.

The unoccupied Trp binding sites in AbeH/FAD/Cl^-^ and apo-AbeH superimpose perfectly with the unoccupied Trp binding site in PyrH/FAD (2WETC). When these structures are compared to PyrH/Trp(2WEU/A), it can be seen that closing of the flavin binding loop in AbeH/FAD/Cl^-^ causes the Glu46 switch to flip pushing Phe49 and Ser50 into the Trp binding pocket by a backbone translation of ∼3 Å and side chain rotation, obstructing the space occupied by the Trp β and α carbons and carboxylate in the PyrH/Trp structure (Figure S14).

The Trp binding site lid, formed by the loop between 3_10_ helix 3 and strand β9 (148-166), is ordered from 152-166 in PyrH/Trp and contains Gln160 and Gln164, both of which form H-bonds with the carboxylate of Trp(30). In both AbeH structures, this loop is disordered from 148- 160, and 161-166 have moved away from the Trp binding site, with the Gln164 side chain amide approximately 8 Å away from its position in PyrH/Trp and facing away from the binding site. Structures of PyrH without Trp bound have the same conformation of the 161-166 portion of the lid as the two AbeH structures.

A simple model for negative coupling in AbeH would suggest that binding of FAD or FADH_2_ to leads to closing of the flavin binding loop, flipping of Glu46 from in to out, and closing the Trp gate by pushing Ser49/Phe50 into the binding site, displacing Trp (Figure S14). The structures of AbeH/FAD and PyrH/FAD (flavin binding loop closed, Glu out, Trp gate closed, lid open) and PyrH/Trp (flavin binding loop open, Glu in, Trp gate open, lid closed) are consistent with this mechanism. Three Trp-6-FDH crystal structures from this clade without Trp bound are consistent with this model (Table S3): Tar14/FAD (6NSD(31), 68%), SttH/FAD/Cl^-^(5HY5(36), 68% identical to AbeH), and apo-Th-Hal (5LV9(29), 67%). FAD-bound SttH and Tar14 have closed flavin binding loops, Glu out, and closed gates, while apo-Th-Hal has an open flavin loop, Glu in, and open gate (29, 31). Tar14, which crystallized in the presence of FAD and Trp, has FAD bound (but not Trp) has a closed flavin binding loop, Glu out, and closed gate, but the lid is closed (30, 31).

There are counterexamples to this simple model. In apo-AbeH, even though the flavin binding loop is open or disordered and Glu46 is pointed inward, the Trp gate is closed, and the lid is open. The closed Trp gate explains our inability to soak Trp into this crystal form, but it is unclear why the gate remains closed even though the flavin binding loop is open. A PyrH/FAD/Trp(30) ternary complex was observed in 2WET-B, which has a closed flavin binding loop with an open gate and closed lid, accommodating both FAD and Trp. The electron density for the isoalloxazine and ribityl groups of FAD is much weaker than the adenosine in this chain, consistent with the fact that full occupancy of the isoalloxazine requires a closed Trp gate.

The residues corresponding to Phe/Ser of the Trp gate in the BorH clade are Val52/Pro53 (BorH numbering), and a similar mechanism coupling flavin binding loop closure and Glu49 switch flipping to blockage of the Trp binding site by the Val52/Pro53 gate has been proposed for Thal. Val/Pro are less bulky than Phe/Ser and the displacement of these residues into the Trp binding site in Thal are smaller, moving about 1 Å rather than 3 Å in the AbeH/PyrH clade. This smaller conformational change could be why FAD binding can eject Trp from AbeH/Trp but not from BorH/Trp, as the conformational change affecting the Trp binding site is apparently greater in the AbeH/PyrH clade. Our BorH structures do not contribute much to our understanding of negative coupling in this clade, as they all have the flavin binding loop disordered and the Val52/Pro53 gate closed, regardless of whether FAD is bound or not because they were all crystallized as apo-BorH and had ligands-soaked in. In our BorH structures as well as published Thal(13, 33) structures, the lid of the Trp binding site has weaker density or is completely disordered when the substrate binding site is vacant.

The partitioning of FAD into BorH chains C and D and Trp into BorH chains A and B can be explained by crystal packing. There is an A/B crystal contact between dimers in adjacent ASUs that sterically hinders access to the FAD binding site is incompatible with the canonically open and closed flavin binding loop conformations and would lead to steric clashes between the loop and FAD in both chains (Figure S15). Since crystals were grown of apo-BorH and then soaked with FAD and Trp into the crystals, this contact prevented binding of FAD to chains A and B. The C/D interface between dimers in the asymmetric unit also involves interactions between the face of BorH containing the FAD binding site, but all three conformations of the flavin loop seen in Thal structures (open, loose, closed) can be modeled at this interface with no steric clashes (Figure S15). However, this close contact between chains C and D still apparently affects the dynamics of the flavin binding loop, as evidenced by the lack of density for the loop and the weaker density for the isoalloxazine in chains C and D. This suggests that the flavin binding loop is not staying “latched” and remains mobile, allowing movement of the isoalloxazine and ribityl groups in and out of the binding groove) while the AMP portion stays fixed. The density following the missing flavin binding loop allows the Glu49 switch to be modeled equally well in both the in and out configurations in BorH/FAD. The fact that the Trp gate (Val52/Pro53) remains in the open position in the FAD bound chains is consistent with the increased mobility of the isoalloxazine.

Since the Trp gate is open in chains C and D, it is not completely clear why Trp is prevented from binding to the FAD-bound chains, (chains C and D). Superposition of those chains with the Trp bound chains (A and B) show no major differences in the position of the Trp gate residues, though the density is weaker and noisier for residues in the Trp lid in chains C and D. There are some slight positional differences between aromatic residues lining the Trp binding site in chains A/B and C/D (Tyr453, Tyr454, Trp465, Phe464), slightly shrinking the Trp binding site in the FAD-bound chains. A network of side chains postulated to be involved in the negative cooperativity in Thal (Tyr362, Asn54, Glu360 in BorH) also show some small positional shifts when comparing the BorH/FAD chains with the BorH/Trp chains, but there are no differences in the hydrogen bonding. Another candidate for a conduit between the flavin and Trp binding sites is an absolutely conserved serine (Ser356 in AbeH, Ser358 in BorH) located approximately equidistant between the FAD and Trp binding sites and which hydrogen bonds with the catalytic lysine. This Ser adopts a different rotamer in the FAD-bound chains than the Trp-bound or apo chains.

BorH and AbeH both have higher affinity for FADH_2_ than for FAD (BorH/FAD *K_D_ =* 13 μM, BorH/FADH_2_ = 0.21 μM, AbeH/FAD *K_D_ =* 1.2 μM, AbeH/FADH_2_ = 0.12 μM), while their flavin reductase counterparts BorF and AbeF have higher affinity for FAD than FADH_2_. (BorF/FAD *K_D_ =* 0.1 μM, BorF/FADH_2_ = 0.8 μM, AbeF/FAD *K_D_ =* 0.8 μM, AbeF/FADH_2_ = 1.5 μM)(23). This is consistent with the directionality of the catalytic cycle (FADH_2_ is the substrate for BorH/AbeH, while FAD is the substrate for BorF/AbeF) and may promote efficient flavin transfer between subunits as has been suggested for other two-component diffusible flavin oxidoreductase systems(39).

Our data suggest that the negative coupling previously observed in Thal is present not only in the closely related BorH but also the more distantly related AbeH and may be an evolutionarily conserved property. The negative coupling likely limits catalytic efficiency, since having one of the two binding sites (flavin or substrate) unoccupied at any given time means that the tunnel connecting the two sites will always be open to bulk solvent at one end or the other, allowing leakage of HOCl/HOBr. HOBr leakage has been previously shown for Thal (11) and tunnel modification has been shown to improve retention of HOBr(37). In the same way, if the structural mechanism for negative coupling could be more completely elucidated, it may be possible to engineer mutations that would “break” the coupling and allow Trp to bind without dissociation of FAD, potentially protecting HOCl/HOBr from solvent and allowing reactions to be carried out at higher flavin concentrations without interfering with Trp binding.

## Conclusion

AbeH is a flavin dependent halogenase which regioselectively chlorinates and brominates Trp at C5 as well as other substrates with indole, phenyl, or quinoline groups. Binding studies indicate that both AbeH and BorH bind FADH_2_ more tightly than FAD and bind Trp with very low affinity. The results of crystallographic and binding studies are consistent with an inability for either AbeH or BorH to form ternary complexes with both FAD/FADH_2_ and Trp. Coupling of the closing of the flavin binding loop and the consequent flipping of the Glu49 switch and closing of the Trp gate explain the inability of FAD and Trp to form a ternary complex with AbeH and also provides a structural explanation by which FAD binding can displace Trp from AbeH/Trp in an entropy-enthalpy compensated manner as our ITC and fluorescence data suggests. If BorH operates like its close homolog Thal, the smaller movement of the Trp gate previously observed in Thal structures upon flavin binding loop closing could be the explanation for why FAD did not seem to displace Trp from BorH/Trp in our ITC experiments, along with the higher Trp and lower FAD affinity observed in our binding studies. However, our partitioned BorH/FAD + BorH/Trp crystal structure could not provide a clear structural explanation, as none of the chains exhibited closing of the flavin binding loop or the Trp gate. A more complete description of the coupling will require a BorH structure with FAD bound and the flavin binding loop closed.

### Experimental Procedures

#### Protein expression and purification

The *abeH* gene was amplified by PCR from a cosmid containing the BE-54017 gene cluster (generously provided by Sean Brady, Rockefeller University) and inserted into expression vector pLIC-HTA using ligation-independent cloning to create an expression construct with an N-terminal His_6_-tag and a TEV protease cleavage site. AbeH was overexpressed in Rosetta (DE3) *E. coli* competent cells (Novagen) grown in Terrific Broth (RPI Corporation) supplemented with 0.1 mg/L ampicillin and 30 mg/mL chloramphenicol and protein expression were induced with 0.1 mM IPTG (isopropyl β-D-thiogalactopyranoside; Gold Biosciences) at 16 °C. Cells were harvested 17–20-hour post-induction. Cell pellets were resuspended in lysis buffer (50 mM Tris-HCl pH 8.2, 500 mM NaCl and 10 mM imidazole) and subjected to three freeze-thaw cycles followed by a 1–2-minute sonication before clarification by centrifugation at 4000 or 12000 rpm. AbeH was captured from clarified lysates using immobilized metal affinity chromatography and further purified by anion exchange and size exclusion chromatography. Purified AbeH in 20 mM HEPES (pH 7.2) and 35 mM sodium citrate was concentrated to 5 mg/mL, flash-frozen, and stored at -80 °C. Protein concentrations were determined by absorbance at 280 nm with calculated extinction coefficients. Protein purity and homogeneity was assessed by SDS-PAGE and gel filtration chromatography (Superdex 200 16/60). BorH, BorF, and AbeF were purified as previously described (23, 24). Proteins used for ITC binding experiments with FADH_2_ were purified in the absence of halide using 20 mM HEPES pH 7.5 100 mM sodium citrate instead of Tris-HCl/NaCl in lysis, affinity chromatography, and gel filtration chromatography buffers.

#### In vitro halogenation of Trp by AbeH

*In vitro* halogenation reactions contained 7 μM AbeH, 13 μM holo-AbeF (copurified with bound FAD), 2.5 mM NADH, 0.5 mM Trp, 50 mM NaCl or NaBr in 10 mM Tris-HCl pH 8.2 in 100 μL reactions. Reactions were initiated by addition of NADH and incubated with 200 rpm shaking at 37 °C. Reactions were stopped by heating to 95 °C for 10 minutes followed by centrifugation at 10,000 x g for 5 minutes. Reaction mixtures were analyzed by reversed phase HPLC using a Kromasil C18 RP-analytical column (100 Å pore size, 4.6 x 250 mm) mounted on a Waters 2695 HPLC. Separation used a 14 minute gradient from 5% B to 100% B (Solvent A= H_2_O/CH_3_CN 95:5, 0.1% TFA; Solvent B = H_2_O/CH_3_CN 5/95, 0.1% TFA) with flow rate 0.6 mL/min. Trp, 5-Cl-Trp, and 5-Br-Trp peaks were monitored with absorbance at 280 nm and retention times and yields were analyzed with using retention times and peak areas from standard curves of authentic standards of Trp (ACROS Organics), 5-Cl-Trp (Santa Cruz Biotechnology) and 5-Br-Trp (Santa Cruz Biotechnology). RP-HPLC purified 5-Cl-Trp and 5-Br-Trp peaks from *in vitro* halogenation reactions were analyzed on a Waters SYNAPT HDMS mass spectrometer using positive ion-mode ESI-MS with leucine enkephalin as internal standard.

To determine the Cl-Trp regioisomer produced by AbeH, 2 mg Trp was chlorinated by incubating 7 μM AbeH, 13 μM holo-AbeF, 2.5 mM NADH, 500 μM Trp, 100 mM NaCl, 10 mM Tris-HCl pH 8.2 for 20 minutes at 37 °C, after which the reaction was stopped by heating to 95 °C for 10 minutes and precipitated proteins were pelleted by centrifugation at 10,000 x g for 5 minutes. Clarified reaction mixtures were concentrated on a silica gel flash column before RP-HPLC purification. The product was dried by rotary evaporation and lyophilization and dissolved in D_2_O for ^1^H NMR analysis using a 600 MHz Bruker Avance III.

Time course experiments were done using 1.7 μM AbeH, 3.3 μM holo-AbeF, 500 μM Trp, 2.5 mM NADH in 10 mM Na/KPO_4_ pH 8.2 with either 50 mM NaCl or 100 mM NaBr. Reactions were incubated for 5 minutes at 37 °C and NADH was added to start the reactions. Aliquots of 100 µL were quenched and analyzed by RP-HPLC at various time points. Unreacted substrate and product concentrations were calculated using peak areas and standard curves of Trp, 5-Cl-Trp, and 5-Br-Trp.

#### Halogenation of non-Trp substrates by AbeH and BorH

To test the ability of BorH and AbeH to chlorinate and brominate other substrates, 100 µL reactions containing 5 µM AbeH or BorH, 10 µM of holo-AbeF or holo-BorF, 100 mM NaCl or NaBr, 2.5 mM NADH, and 0.5 mM substrate in HEPES pH 8.0 were incubated for 12 hours at 37 °C with shaking at 230 rpm. Substrates tested were indole, 5-cyanoindole, serotonin, 5-hydroxytryptophan, tryptamine, 3-indolepropionic acid (3-IPA), anthranilamide, 6-aminoquinoline, 7-aminoquinoline, and 6-amino-2H-1,4-benzoxazin-3(4H)-one. The reactions were stopped by heating at 95 °C for 5 minutes and centrifuged at 12,000 g for 5 minutes. Clarified reaction mixtures were analyzed using RP-HPLC on a Kromasil C18 column (100 Å pore size, 4.6 x 250 mm) mounted on a Waters 2695 HPLC. The gradient was as follows: 0-7.5 min, 5 %-50% B; 7.5-14 min, 50-100 %; 14-25 min, 100 % B; 25-40 min 100-5 % B with flow rate 0.6 mL/min (Solvent A= H_2_O/CH_3_CN 95:5, 0.1% TFA; Solvent B= H_2_O/CH_3_CN 5/95, 0.1% TFA). Substrates and products were detected by monitoring UV absorbance at the *λ*_max_ for each substrate. Product peak fractions were analyzed using ESI-MS on a SYNAPT HDMS mass spectrometer in positive ion mode with leucine enkephalin internal standard.

#### Crystallization and structure determination

AbeH/FAD complex for crystallization was prepared by incubating 50 µM AbeH (in 20 mM HEPES pH 7.2 and 35 mM sodium citrate with 2.5 mM FAD for 20 hours at 4 °C. AbeH/FAD crystals grew overnight in drops containing a 1:1 ratio of AbeH/FAD with reservoir solution (0.1 M Bis-Tris pH 6.2, 0.2 M magnesium acetate, 11% (v/v) PEG 10,000) equilibrated against 1 mL reservoir solution. AbeH/FAD crystals were cryoprotected using reservoir solution supplemented with 20% (v/v) glycerol and flash-frozen in liquid nitrogen for X-ray data collection. To prepare AbeH/FAD/Trp ternary complex crystals by soaking, some AbeH/FAD co-crystals were soaked with Trp (2.5-50 mM) for variable lengths of time (5 minutes to 84 hours.) before cryoprotection and freezing.

Apo-AbeH crystals were grown using the hanging drop vapor diffusion method at room temperature with drops containing 2 μL of protein (40 μM AbeH/20 mM HEPES pH 7.5/350 mM NaCl) and 2 μL of reservoir solution (100 mM Bis-tris propane pH 7.0, 150 mM MgSO_4_, 25% (v/v) PEG 3350) equilibrated against 500 μL of reservoir solution. Apo-AbeH crystals were cryoprotected in reservoir solution containing 20 % (v/v) 2,3-butanediol for 20 minutes, and flash-frozen in liquid nitrogen for X-ray data collection. To prepare AbeH/ Trp ternary complex crystals by soaking, some apo-AbeH crystals were soaked with reservoir solution containing 20 % (v/v) 2,3-butanediol and 50 mM Trp for 50 minutes before freezing.

AbeH/FAD X-ray diffraction data were collected at the Life Sciences Collaborative Access Team beamline 21-ID-F at the Advanced Photon Source (Argonne National Laboratory) with a Rayonix MX 300 detector using the oscillation method (0.5° per frame) (40). Data processing used iMOSFLM for indexing and integration, and Pointless, Aimless, Scala and Truncate for scaling, merging and structure factor generation(41, 42). Molecular replacement was done using Phaser-MR in *Phenix* using 5LV9 (Th-Hal, 66.1% sequence identity to AbeH) as a search model(29, 43, 44). The initial AbeH model was built using AutoBuild, followed with iterative cycles of manual model building in Coot and refinement (rigid body, XYZ, and group B-factors) with Phenix.Refine until both chains were modeled. The FAD binding site was identified visually and using phenix.Ligandfit. Refinement of the AbeH/FAD complex was carried out using Phenix.Refine (XYZ, individual B-factors, occupancies, and TLS parameters). Model validation was performed routinely using MolProbity (45) after each refinement and comprehensive model validation was done in *Phenix* GUI. Apo-AbeH X-ray diffraction data were collected at beamline 21-ID-G and processed using the same method as for AbeH/FAD. The apo-AbeH crystal structure was solved by Phaser-MR using chain B of the AbeH/FAD model. Refinement and model building was performed as described for the AbeH/FAD structure.

Crystals of BorH were grown, soaked, and cryoprotected as described previously for the BorH/Trp crystal structure(24). For this study, apo-BorH crystals, BorH crystals soaked in 6-Cl-Trp, and BorH crystals soaked in both FAD and Trp were frozen and analyzed. X-ray diffraction data were collected at 100 K on a Dectris Eiger X9M detector using the oscillation method (0.25° per frame) at the Life Sciences Collaborative Access Team beamline 21-ID-D at the Advanced Photon Source (Argonne National Laboratory). Data were processed in the same manner as described for AbeH and structures were determined by molecular replacement using Phaser-MR with chain A of the BorH/Trp structure (6UL2.pdb) as a search model. Refinement and structural validation were carried out as described earlier for AbeH.

#### Analysis of Trp, FAD, and FADH_2_ binding by isothermal titration calorimetry

ITC experiments were performed using a VP-ITC isothermal titration calorimeter (MicroCal) in 20 mM HEPES pH 7.5, 350 mM NaCl, 1 mM EDTA and 15% (*v/v*) glycerol. Solutions of AbeH, BorH, FAD (Chem-Implex), and Trp (Acros Organics) were prepared in the same buffer and degassed under vacuum at 30 °C for 20 minutes before loading in the syringe or cell. To analyze formation of binary complexes, AbeH or BorH was placed inside the cell, and ligand (FAD or Trp) was injected. For ternary complex titration, one ligand was preincubated with the macromolecule for 30 minutes at 30 °C to form binary complexes of AbeH/FAD, AbeH/Trp, BorH/FAD, and BorH/Trp before titrating with the second ligand (Figure S16). To test FADH_2_ binding to AbeH and BorH, FAD was reduced to FADH_2_ using sodium dithionite and stored under N_2_. Formation of FADH_2_ from FAD was verified using absorbance and fluorescence spectrophotometry. ITC experiments were performed with reference power set to 10 µcal/sec and Milli-Q water in the reference cell. All experiments were done with a pre-titration delay of 20 minutes, a 5-minute interval between injections, and a filter period of 2 s in high feedback gain mode. All experiments were performed at 310 rpm to mix the components in cell and titrant. All other VP-ITC instrumental parameters were set to default settings. Heats of dilution for all titrants were measured by titrating ligand into buffer for binary complex titrations, and by titrating ligand into a solution containing the other ligand but no protein for ternary complex titration.

The following titrations were carried out:

1.2 mM FAD (2 µL/1 and 5 µL/19 injections) into 18 µM AbeH (Figure 4A)

0.12 mM FADH_2_ (2 µL/1 and 15 µL/19 injections) into 7 µM AbeH in presence of sodium dithionite (Figure 4B)

190 µM FAD (2 µL/1 and 15 µL/29 injections) into 28 µM AbeH preincubated with 16 mM Trp (Figure 4C)

190 µM FAD (2 µL/1 and 15 µL/29 injections) into 28 µM AbeH preincubated with 28 mM Trp (Figure S8)

0.3 mM FAD (2 µL/1 and 10 µL/29 injections) into 8 µM BorH (Figure 4D)

43 µM FADH_2_ (2 µL/1 and 15 µL/19 injections) into 3 µM BorH in presence of sodium dithionite (Figure 4E)

0.63 mM FAD (2 µL/1 and 10 µL/29 injections) into 10 µM BorH preincubated with 18 mM Trp (Figure 4F)

1.4 mM Trp (3 µL/1 and 10 µL/29 injections) into 43 µM AbeH (Figure S7) 16 mM Trp (2 µL/1 and 10 µL/29 injections) into 18 µM AbeH (Figure 5A)

27 mM Trp (2 µL/1 and 10 µL/29 injections) into 48 µM AbeH preincubated with 3.3 mM FAD (Figure 5B)

0.35 mM Trp (2 µL/1 and 15 µL/19 injections) into 11 µM AbeH preincubated with 0.1 mM FADH_2_ in presence of sodium dithionite (Figure 5C)

21 mM Trp (2 µL/1 and 10 µL/29 injections) into 8 µM BorH (Figure 5D)

21 mM Trp (2 µL/1 and 10 µL/29 injections) into 8 µM BorH preincubated with 5 mM FAD (Figure 5E)

0.35 mM Trp (2 µL/1 and 15 µL/19 injections) into 11 µM BorH preincubated with 0.1 mM FADH_2_ in presence of sodium dithionite (Figure 5F)

Titrations were analyzed by manually integrating each injection, subtracting the heat of dilution, and carrying out non-linear least squares fit of the calorimetric binding data to a one site model using Origin 7.0 (Origin Lab). Trp binding experiments were subtracted by point-by-point heat of dilution, and FAD and FADH_2_ binding experiments were subtracted using the mean heat of the last three injections. For the Trp binding data, *n* was fixed to 1 based on expected stoichiometry of 1:1 based on FDH crystal structures and as proposed for lower c-value titrations in the literature (24, 25, 46). The other parameters were not constrained during fitting.

#### Analysis of Trp and FAD binding by fluorescence spectroscopy

Quenching of FAD fluorescence quenching upon binding to AbeH was measured in the absence and presence of Trp in 20 mM Tris-HCl pH 8.0 and 100 mM NaCl using a Quantmaster 40 fluorimeter (Photon Technologies International). Trp and FAD solutions were prepared in 20 mM Tris-HCl pH 8, and concentrations were determined by absorbance at 280 nm and 450 nm respectively absorbance values. FAD fluorescence was measured using excitation at 450 nm (4 nm slit opening) and emission set at 525 nm (6 nm slit opening) using a fluorescence cuvette to collect 50 scans, which were averaged later during data processing in Microsoft Excel. Experiments were performed in duplicate by adding different concentrations of AbeH into fixed FAD (0.45-0.50 mM) and without or with different fixed Trp (0.3, 5.5 mM) concentrations. Binding experiments were performed by mixing the components and incubating for two minutes before data collection by excitation at 450 nm and emission set at 525 nm. Raw fluorescence intensities were corrected for the buffer, Trp, and AbeH background contributions. Quenched fluorescence intensity was plotted vs. AbeH concentration and fit to equation (1) using GraphPad Prism 8.0 by nonlinear regression analysis. In equation 1, F = fractional saturation, ΔF= fluorescence quenching upon AbeH/FAD complex formation (calculated from the difference of FAD fluorescence intensity at different AbeH concentration), ΔF_max_= maximum fluorescence change obtained from fit, [AbeH_FAD_]= FAD bound AbeH, [AbeH]_T_= Total AbeH concentration, [FAD]_T_= Total FAD concentration, *K_D_* = equilibrium dissociation constant.

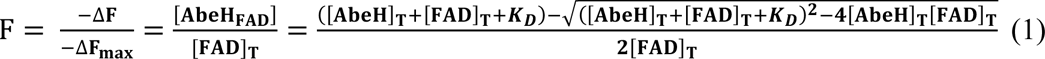

#### Intrinsic tryptophan fluorescence spectroscopy

Intrinsic tryptophan fluorescence spectroscopy was used to measure FAD binding to AbeH and BorH in 20 mM HEPES pH 7.7- and 50-mM sodium citrate. FAD was also prepared in the same buffer and concentration was determined spectrophotometrically as described earlier. Fluorescence experiments were performed with Photon Technologies International using identical instrumental set up described earlier. FAD binding to apo-AbeH or apo-BorH was performed in duplicate with 100 nM fixed AbeH or BorH with variable FAD concentrations up to 30 µM and 70 µM, respectively. AbeH/FAD binding was monitored using excitation at 295 nm and emission at 328 nm. BorH/FAD binding was monitored using excitation at 290 nm and emission at 326 nm. Inner filter effect corrected fluorescence intensities were plotted against quenched intrinsic tryptophan fluorescence of AbeH and BorH. Data were fitted by non-linear regression to equation 1 using GraphPad Prism 8.0.

### Author Contributions

M.A. carried out expression and purification of AbeH, AbeF, and BorH, AbeH halogenase activity assays, crystallization, X-ray diffraction data collection, structure determination, refinement, model building, and structural analysis of apo-AbeH and AbeH/FAD, and the ITC and fluorescence titration experiments on AbeH and BorH. K.L. expressed and purified BorH and BorF and carried out crystallization, X-ray diffraction data collection, molecular replacement, model building, and refinement of apo-BorH, BorH/FAD/Trp, and BorH/6-Cl-Trp. A.J.D.S. carried out expression and purification of AbeH, AbeF, BorH, and BorF, and carried out BorH and AbeH halogenation assays on non-Trp substrates. J.J.B. carried out model building and refinement, data analysis, and structural analysis. The manuscript was written by M.A. and J.J.B. with input from K.L. and A.J.D.S.

### Data Availability

All X-ray crystal structures have had coordinates and structure factors deposited in the RCSB Protein Data Bank with the following accession numbers: 8FOX (Apo-AbeH), 8FOV (AbeH/FAD), 8TTJ (BorH/6-Cl-Trp), 8TTK(Apo-BorH), 8TTI (BorH/FAD+BorH/Trp).

## Abbreviations

FDH: (flavin-dependent halogenase)
FR: (flavin reductase)
ESI-MS: (Electrospray ionization mass spectroscopy)
ITC: (isothermal titration calorimetry)
Trp: (*L*-tryptophan)
5-Cl-Trp: (5-chloro-*L*- tryptophan)
5-Br-Trp: (5-bromo-*L*-tryptophan)
6-Cl-Trp: (6-chloro-*L*-tryptophan)
*t_R_*: (retention time)
*m*/*z*: (mass to charge ratio)
RP-HPLC: (Reversed-phase high-pressure liquid chromatography)
IMAC: (Immobilized metal affinity chromatography) root mean square deviation (rmsd) asymmetric unit (ASU).

## Acknowledgements

The authors wish to thank J. David Dignam and Tim Mueser for expert advice and assistance with binding studies and the Center for Drug Design at the University of Toledo for the use of analytical instrumentation. This research used resources of the Advanced Photon Source; a U.S. Department of Energy (DOE) Office of Science User Facility operated for the DOE Office of Science by Argonne National Laboratory under Contract No. DE-AC02-06CH11357. Use of the LS-CAT Sector 21 was supported by the Michigan Economic Development Corporation and the Michigan Technology Tri-Corridor (Grant 085P1000817). This work was supported by the National Institutes of Health (award 1R15GM144877-01 to J.J.B.) and the University of Toledo Research Awards and Fellowships Program (award 206207 to J.J.B.). The content is solely the responsibility of the authors and does not necessarily represent the official views of the National Institutes of Health.

## Conflict of Interest

The authors declare that they have no conflicts of interest with the contents of this article.

## Notes

### Competing Interest Statement

The authors have declared no competing interest.

## References

1. van Pée, K. H., and Patallo, E. P. (2006) Flavin-dependent halogenases involved in secondary metabolism in bacteria Appl Microbiol Biotechnol 70, 631–641 10.1007/s00253-005-0232-2

2. Neumann, C. S., Fujimori, D. G., and Walsh, C. T. (2008) Halogenation strategies in natural product biosynthesis Chem Biol 15, 99–109 10.1016/j.chembiol.2008.01.006

3. Keller, N. P. (2019) Fungal secondary metabolism: regulation, function and drug discovery Nat Rev Microbiol 17, 167–180 10.1038/s41579-018-0121-1

4. Phintha, A., Prakinee, K., and Chaiyen, P. (2020) Chapter Eleven - Structures, mechanisms and applications of flavin-dependent halogenases In The Enzymes, Chaiyen P, Tamanoi F, eds. Academic Press, 327–364

5. Frese, M., Schnepel, C., Minges, H., Voß, H., Feiner, R., and Sewald, N. (2016) Modular Combination of Enzymatic Halogenation of Tryptophan with Suzuki–Miyaura Cross-Coupling Reactions ChemCatChem 8, 1799–1803 10.1002/cctc.201600317

6. Palani, V., Perea, M. A., and Sarpong, R. (2022) Site-Selective Cross-Coupling of Polyhalogenated Arenes and Heteroarenes with Identical Halogen Groups Chem Rev 122, 10126–10169 10.1021/acs.chemrev.1c00513

7. Schroeder, L., Frese, M., Müller, C., Sewald, N., and Kottke, T. (2018) Photochemically Driven Biocatalysis of Halogenases for the Green Production of Chlorinated Compounds ChemCatChem 10, 3336–3341 10.1002/cctc.201800280

8. Crowe, C., Molyneux, S., Sharma, S. V., Zhang, Y., Gkotsi, D. S., Connaris, H. et al. (2021) Halogenases: a palette of emerging opportunities for synthetic biology–synthetic chemistry and C–H functionalisation Chem Soc Rev 50, 9443–9481 10.1039/D0CS01551B

9. Karabencheva-Christova, T. G., Torras, J., Mulholland, A. J., Lodola, A., and Christov, C. Z. (2017) Mechanistic Insights into the Reaction of Chlorination of Tryptophan Catalyzed by Tryptophan 7-Halogenase Sci Rep 7, 17395 10.1038/s41598-017-17789-x

10. Barker, R. D., Yu, Y., De Maria, L., Johannissen, L. O., and Scrutton, N. S. (2022) Mechanism of Action of Flavin-Dependent Halogenases ACS Catal 12, 15352–15360 10.1021/acscatal.2c05231

11. Phintha, A., Prakinee, K., Jaruwat, A., Lawan, N., Visitsatthawong, S., Kantiwiriyawanitch, C., et al. (2021) Dissecting the low catalytic capability of flavin-dependent halogenases J Biol Chem 296, 10.1074/jbc.RA120.016004

12. Sana, B., Ho, T., Kannan, S., Ke, D., Li, E. H. Y., Seayad, J. et al. (2021) Engineered RebH Halogenase Variants Demonstrating a Specificity Switch from Tryptophan towards Novel Indole Compounds ChemBioChem 22, 2791–2798 10.1002/cbic.202100210

13. Moritzer, A. C., Minges, H., Prior, T., Frese, M., Sewald, N., and Niemann, H.H. (2019) Structure-based switch of regioselectivity in the flavin-dependent tryptophan 6- halogenase Thal J Biol Chem 294, 2529–2542 10.1074/jbc.RA118.005393

14. Glenn, W. S., Nims, E., and O’Connor, S. E. (2011) Reengineering a tryptophan halogenase to preferentially chlorinate a direct alkaloid precursor J Am Chem Soc 133, 19346–19349 10.1021/ja2089348

15. Jiang, Y., and Lewis, J. C. (2023) Asymmetric catalysis by flavin-dependent halogenases Chirality 35, 452–460 10.1002/chir.23550

16. Payne, J. T., Andorfer, M. C., and Lewis, J. C. (2016) Chapter Five - Engineering Flavin-Dependent Halogenases In Meth Enzymol, O’Connor SE, ed. Academic Press, 93–126

17. Dachwitz, S., Widmann, C., Frese, M., Niemann, H. H., and Sewald, N. (2021) Enzymatic halogenation: enzyme mining, mechanisms, and implementation in reaction cascades In Amino Acids, Peptides and Proteins: Volume 44, The Royal Society of Chemistry, 1–43

18. Andorfer, M. C., and Lewis, J. C. (2018) Understanding and Improving the Activity of Flavin-Dependent Halogenases via Random and Targeted Mutagenesis Annu Rev Biochem 87, 159–185 10.1146/annurev-biochem-062917-012042

19. Latham, J., Brandenburger, E., Shepherd, S. A., Menon, B. R. K., and Micklefield, J. (2018) Development of Halogenase Enzymes for Use in Synthesis Chem Rev 118, 232–269 10.1021/acs.chemrev.7b00032

20. Milshteyn, A., Schneider, J. S., and Brady, S. F. (2014) Mining the metabiome: identifying novel natural products from microbial communities Chem Biol 21, 1211–1223 10.1016/j.chembiol.2014.08.006

21. Chang, F. Y., and Brady, S. F. (2011) Cloning and characterization of an environmental DNA-derived gene cluster that encodes the biosynthesis of the antitumor substance BE- 54017 J Am Chem Soc 133, 9996–9999 10.1021/ja2022653

22. Chang, F. Y., and Brady, S. F. (2013) Discovery of indolotryptoline antiproliferative agents by homology-guided metagenomic screening Proc Natl Acad Sci U S A 110, 2478–2483 10.1073/pnas.1218073110

23. De Silva, A. J., Sehgal, R., Kim, J., and Bellizzi, J. J. (2021) Steady-state kinetic analysis of halogenase-supporting flavin reductases BorF and AbeF reveals different kinetic mechanisms Arch Biochem Biophys 704, 108874 10.1016/j.abb.2021.108874

24. Lingkon, K., and Bellizzi, J. J. (2020) Structure and Activity of the Thermophilic Tryptophan-6 Halogenase BorH ChemBioChem 21, 1121–1128 10.1002/cbic.201900667

25. Dong, C., Flecks, S., Unversucht, S., Haupt, C., van Pée, K. H., and Naismith, J. H. (2005) Tryptophan 7-halogenase (PrnA) structure suggests a mechanism for regioselective chlorination Science 309, 2216–2219 10.1126/science.1116510

26. Urzhumtseva, L., Afonine, P. V., Adams, P. D., and Urzhumtsev, A. (2009) Crystallographic model quality at a glance Acta Crystallogr D Biol Crystallogr 65, 297–300 10.1107/S0907444908044296

27. Bitto, E., Huang, Y., Bingman, C. A., Singh, S., Thorson, J. S., and Phillips Jr, G. N. (2008) The structure of flavin-dependent tryptophan 7-halogenase RebH Proteins: Struct Funct Genet 70, 289–293 10.1002/prot.21627

28. Yeh, E., Blasiak, L. C., Koglin, A., Drennan, C. L., and Walsh, C. T. (2007) Chlorination by a long-lived intermediate in the mechanism of flavin-dependent halogenases Biochemistry 46, 1284–1292 10.1021/bi0621213

29. Menon, B. R., Latham, J., Dunstan, M. S., Brandenburger, E., Klemstein, U., Leys, D. et al. (2016) Structure and biocatalytic scope of thermophilic flavin-dependent halogenase and flavin reductase enzymes Org Biomol Chem 14, 9354–9361 10.1039/c6ob01861k

30. Zhu, X., De Laurentis, W., Leang, K., Herrmann, J., Ihlefeld, K., van Pée, K.-H. et al. (2009) Structural Insights into Regioselectivity in the Enzymatic Chlorination of Tryptophan J Mol Biol 391, 74–85 10.1016/j.jmb.2009.06.008

31. Luhavaya, H., Sigrist, R., Chekan, J. R., McKinnie, S. M. K., and Moore, B. S. (2019) Biosynthesis of l-4-Chlorokynurenine, an Antidepressant Prodrug and a Non-Proteinogenic Amino Acid Found in Lipopeptide Antibiotics Angew Chem Int Ed 58, 8394–8399 10.1002/anie.201901571

32. Widmann, C., Ismail, M., Sewald, N., and Niemann, H. H. (2020) Structure of apo flavin-dependent halogenase Xcc4156 hints at a reason for cofactor-soaking difficulties Acta crystallogr D Biol Crystallogr 76, 687–697 10.1107/s2059798320007731

33. Moritzer, A. C., and Niemann, H. H. (2019) Binding of FAD and tryptophan to the tryptophan 6-halogenase Thal is negatively coupled Prot Sci 28, 2112–2118 10.1002/pro.3739

34. Velazquez-Campoy, A., Goñi, G., Peregrina, J. R., and Medina, M. (2006) Exact analysis of heterotropic interactions in proteins: Characterization of cooperative ligand binding by isothermal titration calorimetry Biophys J 91, 1887–1904 10.1529/biophysj.106.086561

35. Brown, A. (2009) Analysis of cooperativity by isothermal titration calorimetry Int J Mol Sci 10, 3457–3477 10.3390/ijms10083457

36. Shepherd, S. A., Menon, B. R. K., Fisk, H., Struck, A.-W., Levy, C., Leys, D. et al. (2016) A Structure-Guided Switch in the Regioselectivity of a Tryptophan Halogenase ChemBioChem 17, 821–824 10.1002/cbic.201600051

37. Prakinee, K., Phintha, A., Visitsatthawong, S., Lawan, N., Sucharitakul, J., Kantiwiriyawanitch, C. et al. (2022) Mechanism-guided tunnel engineering to increase the efficiency of a flavin-dependent halogenase Nat Catal 5, 534–544 10.1038/s41929-022-00800-8

38. Jeon, J., Lee, J., Jung, S.-M., Shin, J. H., Song, W. J., and Rho, M. (2021) Genomic Determinants Encode the Reactivity and Regioselectivity of Flavin-Dependent Halogenases in Bacterial Genomes and Metagenomes mSystems 6, 10.1128/msystems.00053-00021 10.1128/msystems.00053-21

39. Sucharitakul, J., Tinikul, R., and Chaiyen, P. (2014) Mechanisms of reduced flavin transfer in the two-component flavin-dependent monooxygenases Arch Biochem Biophys 555-556, 33–46 10.1016/j.abb.2014.05.009

40. Otwinowski, Z., and Minor, W. (1997) Processing of X-ray diffraction data collected in oscillation mode Methods Enzymol 276, 307–326 10.1016/s0076-6879(97)76066-x

41. Battye, T. G., Kontogiannis, L., Johnson, O., Powell, H. R., and Leslie, A. G. (2011) iMOSFLM: a new graphical interface for diffraction-image processing with MOSFLM Acta Crystallogr D Biol Crystallogr 67, 271–281 10.1107/s0907444910048675

42. Potterton, E., McNicholas, S., Krissinel, E., Cowtan, K., and Noble, M. (2002) The CCP4 molecular-graphics project Acta Crystallogr D Biol Crystallogr 58, 1955–1957 10.1107/s0907444902015391

43. Adams, P. D., Afonine, P. V., Bunkóczi, G., Chen, V. B., Davis, I. W., Echols, N. et al. (2010) PHENIX: a comprehensive Python-based system for macromolecular structure solution Acta Crystallogr D Biol Crystallogr 66, 213–221 10.1107/s0907444909052925

44. Adams, P. D., Afonine, P. V., Bunkóczi, G., Chen, V. B., Davis, I. W., Echols, N. et al. (2010) PHENIX: a comprehensive Python-based system for macromolecular structure solution Acta Crystallogr D Biol Crystallogr 66, 213–221 10.1107/s0907444909052925

45. Williams, C. J., Headd, J. J., Moriarty, N. W., Prisant, M. G., Videau, L. L., Deis, L. N. et al. (2018) MolProbity: More and better reference data for improved all-atom structure validation Prot Sci 27, 293–315 10.1002/pro.3330

46. Turnbull, W. B., and Daranas, A. H. (2003) On the value of c: can low affinity systems be studied by isothermal titration calorimetry? J Am Chem Soc 125, 14859–14866 10.1021/ja036166s

