## Supporting Information for "Crystallographic and thermodynamic evidence of negative cooperativity of flavin and tryptophan binding in the flavin-dependent halogenases AbeH and BorH"

Figure S1: Purification of AbeH

Figure S2: AbeH chlorinates and brominates Trp *in vitro*

Table S1: Mass spectrometry data for halogenated products produced by BorH and AbeH

Figure S3: $^1$H NMR of Trp chlorination product of AbeH

Figure S4: Time-course for *in vitro* Trp halogenation by AbeH

Figure S5: AbeH chlorinates and brominates non-tryptophan aromatic substrates

Figure S6. BorH chlorinates and brominates non-tryptophan aromatic substrates

Figure S7: No binding detected by ITC when Trp is titrated into AbeH at low molar ratios.

Figure S8: Confirmation of endothermic titration of FAD into AbeH/Trp

Figure S9: Quenching of intrinsic Trp fluorescence of AbeH and BorH upon titration with FAD

Figure S10: Quenching of FAD fluorescence upon titration with AbeH or BorH

Table S2: ITC determined thermodynamic parameters of FAD, $FADH_2$, and Trp binding to AbeH and BorH

Figure S11: Phylogenetic tree of AbeH and BorH with other FDHs with solved crystal structures

Figure S12: Multiple sequence alignment of AbeH and BorH with other FDHs

Figure S13: Difference in Trp binding lid between BorH clade and AbeH clade with solved crystal structures

Figure S14: Movement of the Trp gate upon FAD binding creates steric hindrance for Trp

Figure S15: Crystal packing prevents soaking of FAD into chains A and B of BorH/Trp + BorH/FAD

Table S3. Conformations of the flavin binding strap loop and Trp binding site observed in crystal structures of FDHs in the AbeH clade

Figure S16: ITC experiments designed to test ternary complex formation in FDH

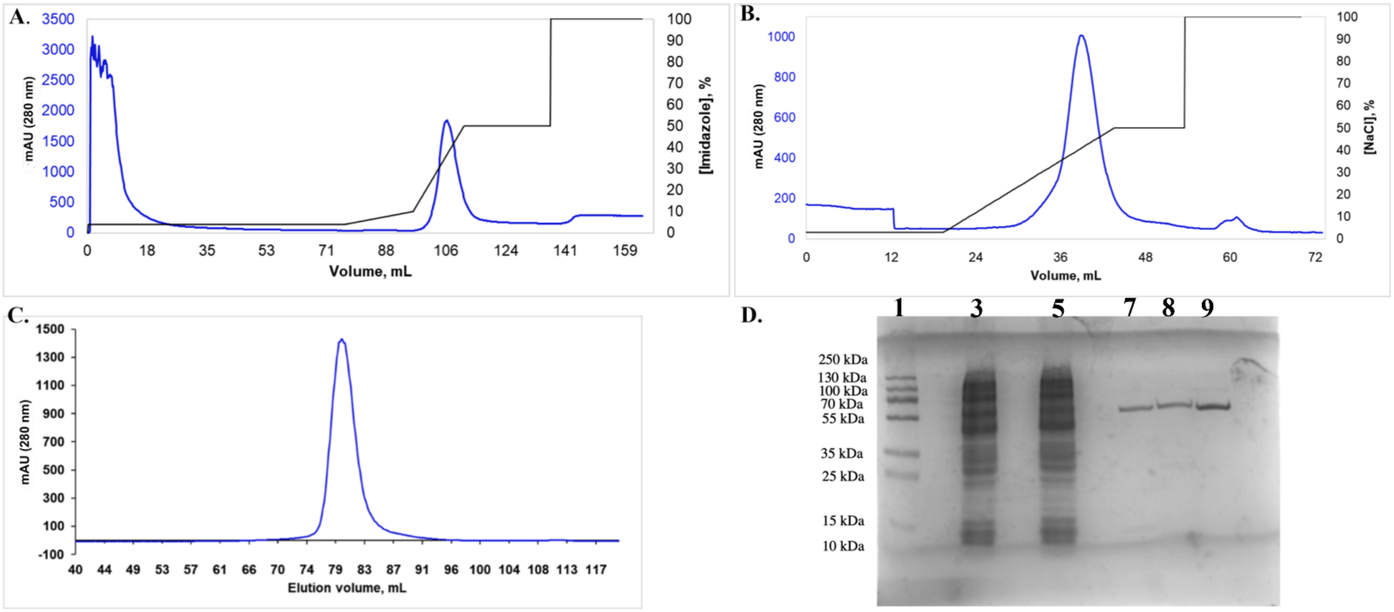


**Figure S1: Purification of AbeH.** All steps were carried out at 4 ºC on an Akta Purifier 10 FPLC. Data was analyzed on Unicorn 5.0 and chromatograms were plotted using Microsoft Excel.

**A.** Purification of AbeH by immobilized metal ion affinity chromatography (IMAC). AbeH was lysed in 50 mM Tris-HCl pH 8.2, 0.5 M NaCl, and 10 mM imidazole and purified on a 5 mL $Co^{2+}$-charged HisTrap crude FF column (GE Biosciences) in 50 mM Tris-HCl pH 8.2, 100 mM NaCl with a gradient from 10-250 mM imidazole. $A_{280}$ trace is in blue, and imidazole gradient is in black. AbeH elutes at ~100 mM imidazole. Pooled peak fractions from this column are shown on lane 7 on the gel in **D**.

**B.** Purification of AbeH by anion exchange chromatography. Pooled IMAC fractions were diluted three-fold with 20 mM Tris-HCl pH 8.2 and loaded on a 5 mL HiTrap Q FF column and purified with a gradient of 0-1 M NaCl in 20 mM Tris-HCl pH 8.2. $A_{280}$ trace is in blue, and NaCl gradient is in black. AbeH elutes at 400 mM NaCl. Pooled peak fractions from this column are shown on lane 8 on the gel in **D**.

**C.** Purification of AbeH by gel filtration chromatography. Pooled anion exchange fractions were concentrated to 2 mL and injected on a Superdex 200 16/60 size exclusion column and purified in 20 mM HEPES pH 7.2- and 35-mM sodium citrate. AbeH elutes at 80 mL, corresponding to an apparent solution molecular mass of 94 kDa based on a gel filtration standard curve. Pooled peak fractions from this column are shown on lane 9 on the gel in **D**.

**D.** 12% SDS-PAGE of AbeH purification stained with Coomassie blue. Lane 1: Molecular weight ladder; Lane 3: Soluble lysate; Lane 5: IMAC flow-through. Lane 7: Pooled IMAC fractions. Lane 8: Pooled anion exchange fractions. Lane 9: Pooled gel filtration fractions. AbeH with N-terminal $His_6$-tag has a molecular weight of 59.5 kDa.

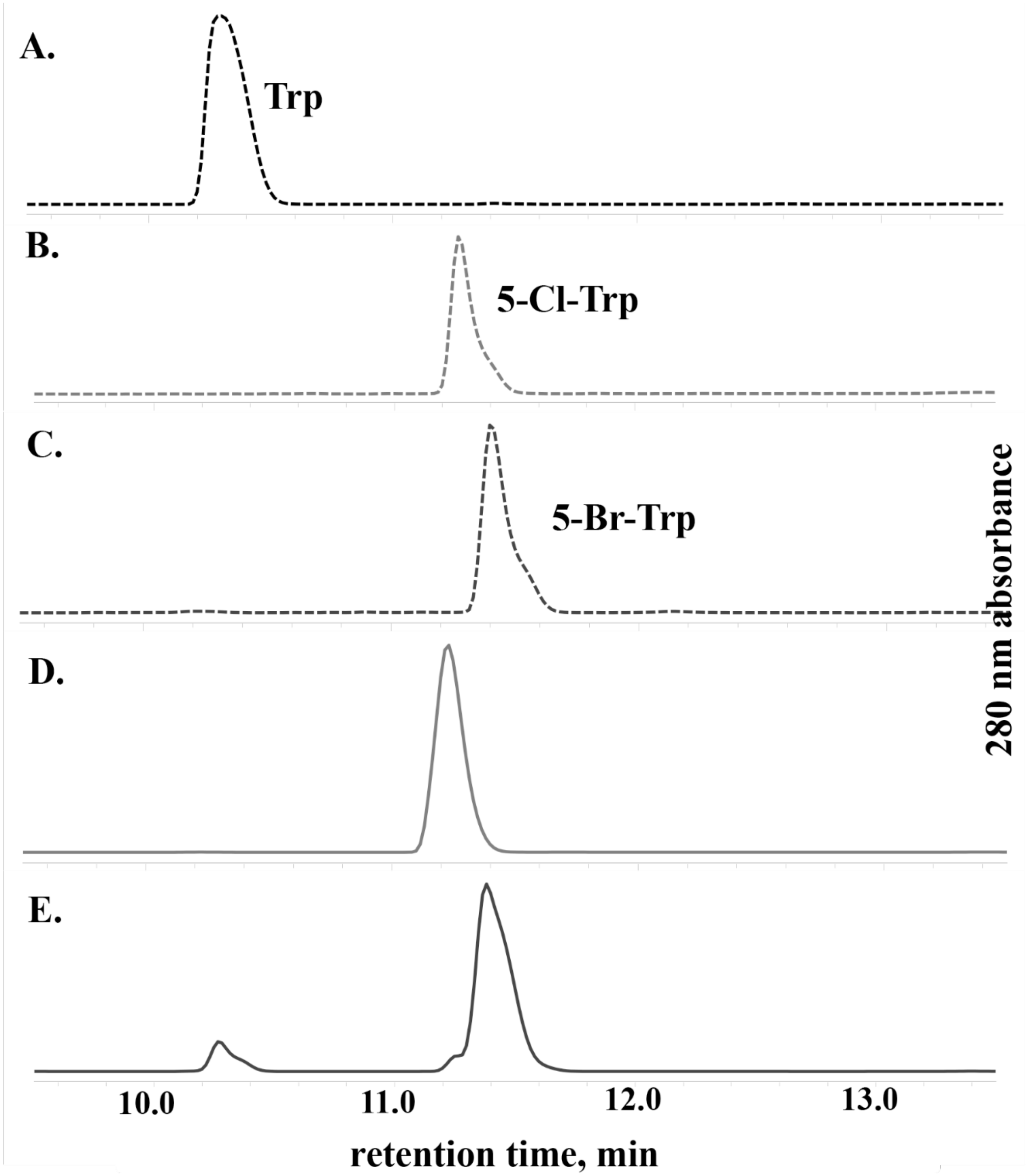


**Figure S2: AbeH chlorinates and brominates Trp *in vitro*.**

*In vitro* halogenation reactions were performed with 7 μM AbeH, 13 μM holo-AbeF, 2.5 mM NADH, 50 mM NaCl or NaBr in 10 mM Tris-HCl pH 8.2 at 37 °C shaking at 200 rpm with final volume of 100 μL. Standards and reactions were analyzed on a Kromasil C18 column (100 Å pore size, 4.6 x 250 mm) mounted on a Waters HPLC using a 14 min gradient from 5% -100% B (Solvent A:5% $CH_3CN$+0.1% TFA; Solvent B:95% $CH_3CN$+0.1% TFA) with flow rate 0.6 mL/min using 90 μL injections. Baseline subtracted peak integration was performed on Empower software (Waters). **(A)** Trp standard ($t_R$ = 10.3 min). **(B)** 5-Cl-Trp standard ($t_R$ =11.3 min). **(C)** 5-Br-Trp standard ($t_R$ = 11.4 min). **(D)** 60 min AbeH *in vitro* chlorination reaction of 0.5 mM Trp ($t_R$ = 11.3 min, 100% conversion to 5-Cl-Trp). **(E)** 60 min AbeH *in vitro* bromination reaction of 0.5 mM Trp ($t_R$ = 10.3 min and 11.5 min, ~80% conversion to 5-Br-Trp).

**Table S1: Mass spectrometry data for halogenated products produced by BorH and AbeH.**

| Substrate | MW of monochlorinated substrate (g/mol) | MW of monobrominated substrate (g/mol) | BorH | | AbeH | |
|---|---|---|---|---|---|---|
| | | | *m/z* of product peaks from NaCl reaction (intensity ratio) | *m/z* of product peaks from NaBr reaction (intensity ratio) | *m/z* of product peaks from NaCl reaction (intensity ratio) | *m/z* of product peaks from NaBr reaction (intensity ratio) |
| tryptophan | 238.67 | 283.12 | 239.0612, 241.0559 (3:1) | 283.0103, 285.0078 (1:1) | 239.0618, 241.0627 (3:1) | 283.0109, 285.0478 (1:1) |
| indole | 151.59 | 196.04 | 152.0784, 153.9882 (3.1:1) | 195.1597, 197.1432 (1:1) | 152.0832, 154.0231 (2.8:1) | 195. 1632, 197.0873 (1.2:1) |
| 5-cyanoindole | 176.60 | 221.05 | 177.6148, 179.7031 (2.9:1) | 222.0642, 224.1042 (1.1:1) | No activity | |
| serotonin | 210.65 | 255.10 | 211.0622, 213.0731 (3:1) | 256.0953, 258.1032 (1:1) | No activity | |
| 5-hydroxytryptophan | 254.66 | 299.11 | 255.0534, 257.0523 (3:1) | 299.0053, 301.0432 (1.2:1) | No activity | |
| tryptamine | 194.66 | 239.11 | 195.0364, 197.0634 (2.8:1) | 239.0234, 241.0324 (1:1) | No activity | |
| 3-indolepropionic acid | 223.65 | 268.10 | 224.0367, 226.0894 (2.9:1) | 267.9931, 269.9993 (1:1.1) | 224.0597, 226.1264 (2.6:1) | 267.9922, 269.9944 (1.2:1) |
| anthranilamide | 170.59 | 215.04 | 171.0326, 173.0303 (2.7:1) | 214.9831, 216.9678 (1:1.2) | 171.1539, 173.0298 (3:1) | 214.1039, 216.9925 (1:1.1) |
| 6-aminoquinoline | 178.61 | 223.06 | 179.0372, 181.9633 (2.8:1) | 222.9869, 224.9842 (1.1:1) | 179.0392, 181.9626 (2.7:1) | 222.9928, 224.9910 (1.1:1) |
| 7-aminoquinoline | 178.61 | 223.06 | No activity | | 179.0298, 181.8945 (2.9:1) | 222.9967, 224.9904 (1:1) |
| 6-amino-2H-1,4-benzoxazin-3(4H)-one | 198.60 | 243.05 | 199.1729, 201.1693 (2.3:1) | 244.0268, 246.0481 (1.1:1) | No activity | |

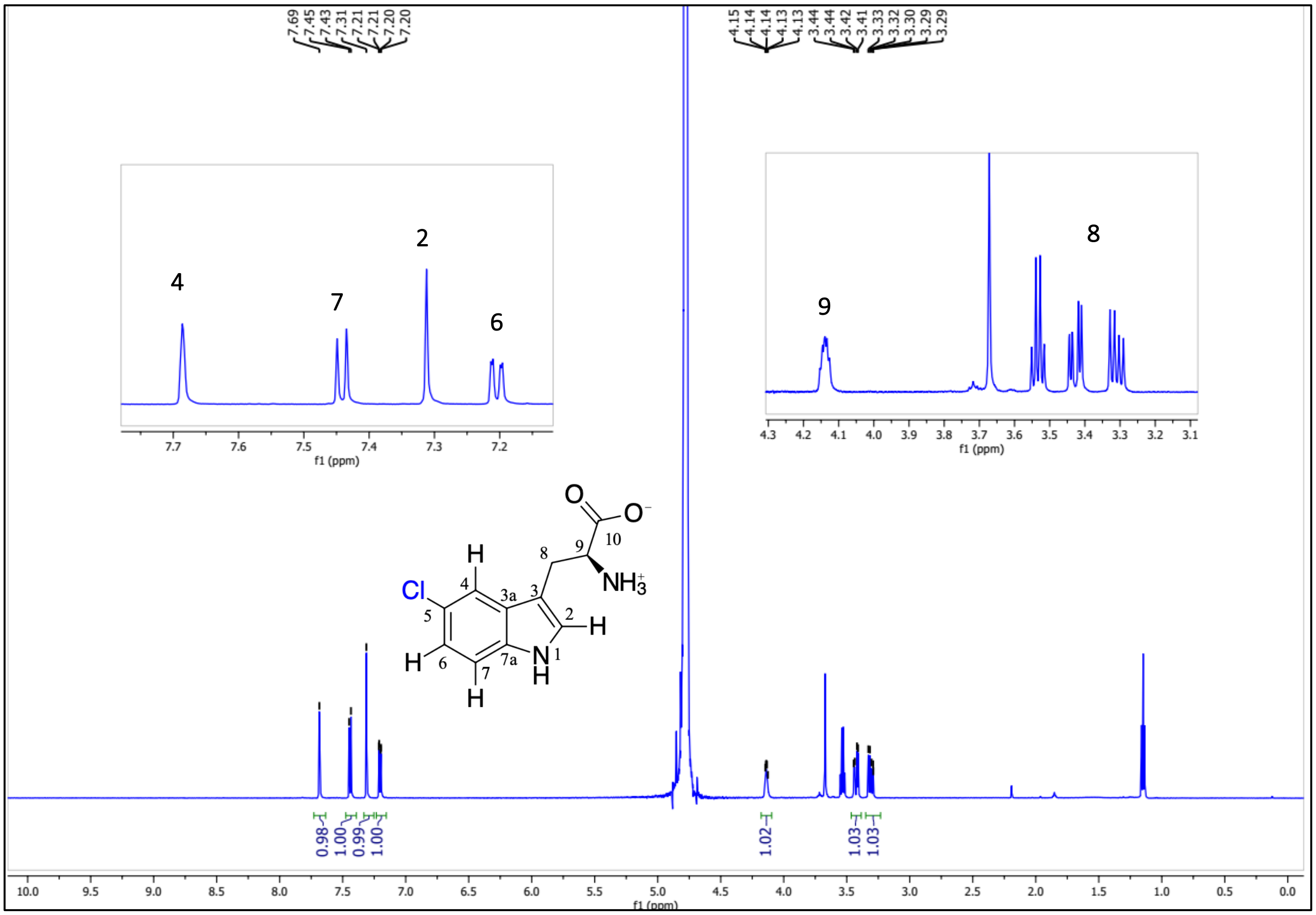

**Figure S3: $^{1}$H NMR spectrum of 5-Cl-Trp produced by *in vitro* chlorination of Trp by AbeH**

$^{1}$H NMR (600 MHz, $D_2O$) spectrum of product isolated by HPLC.: δ 7.69 (s, 1H, ArH4), 7.44 (d, *J* = 8.7 Hz, 1H, ArH7), 7.31 (s, 1H, ArH2), 7.20 (dd, *J* = 8.7, 1.7 Hz, 1H, ArH6), 4.25 – 4.04 (m, 1H, H9), 3.43 (dd, *J* = 15.4, 5.1 Hz, 1H, H8A), 3.31 (dd, *J* = 15.4, 7.6 Hz, 1H, H8B).

**A.** **B.**

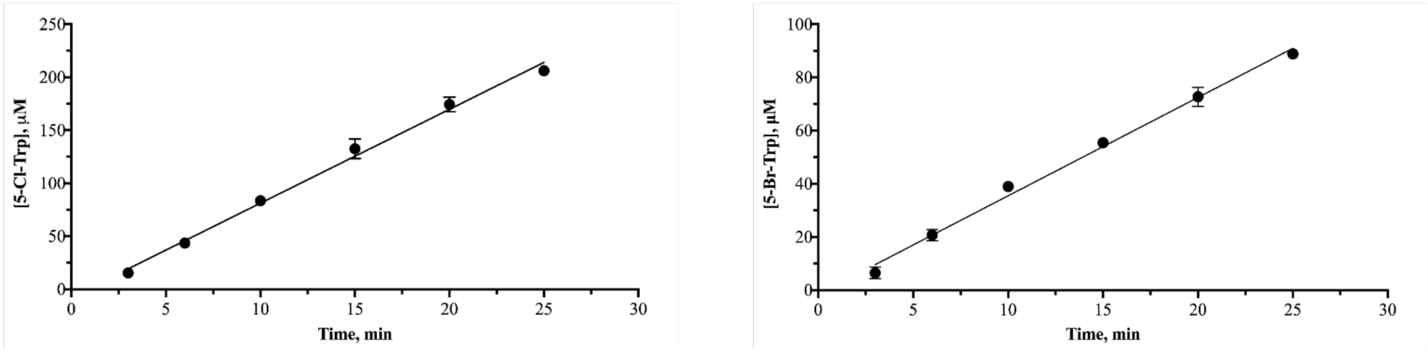


**Figure S4: Time-course of *in vitro* Trp chlorination and bromination by AbeH.**

Reaction conditions: 1.7 µM AbeH, 3.3 µM holo-AbeF, 0.5 mM Trp, 2.5 mM NADH, 10 mM Na/$KPO_4$ pH 8.2, and either 50 mM NaCl (**A**) or 100 mM NaBr (**B**). Concentration of product formed was calculated from RP-HPLC chromatogram integrated $A_{280}$ peak areas compared to standard curves of integrated peak areas of known concentrations of 5-Cl-Trp and 5-Br-Trp standards. Experiments were carried out in duplicate. **(A)** Time course of Trp chlorination by AbeH has chlorination specific activity of 87 µmol $min^{-1}$ $mg^{-1}$. **(B)** Time course of Trp bromination by AbeH has bromination specific activity of 34 µmol $min^{-1}$ $mg^{-1}$.

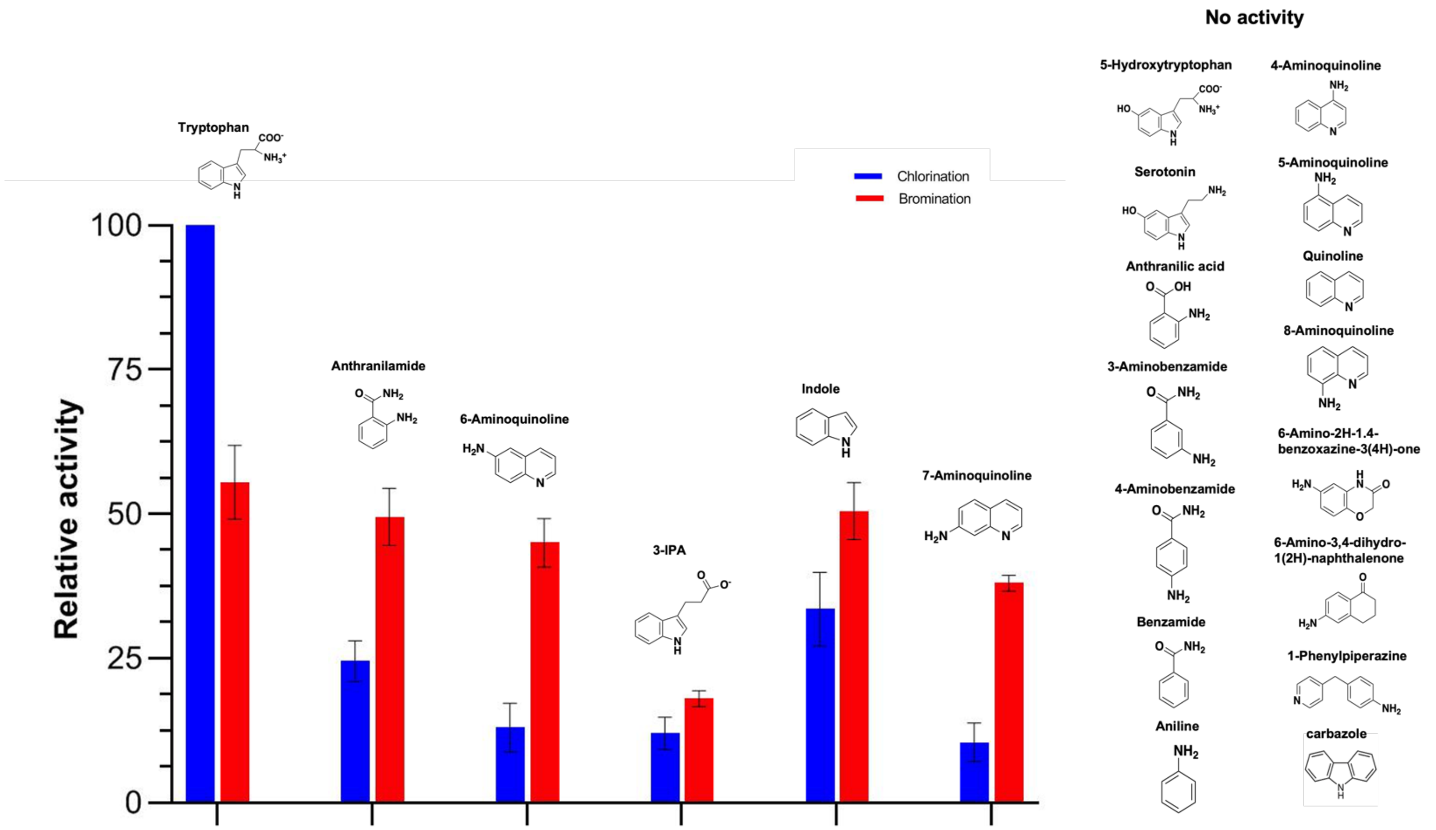


**Figure S5. AbeH chlorinates and brominates non-tryptophan aromatic substrates.**

Results from *in vitro* chlorination and bromination assays of AbeH with Trp and 20 other aromatic substrates under identical reaction conditions.

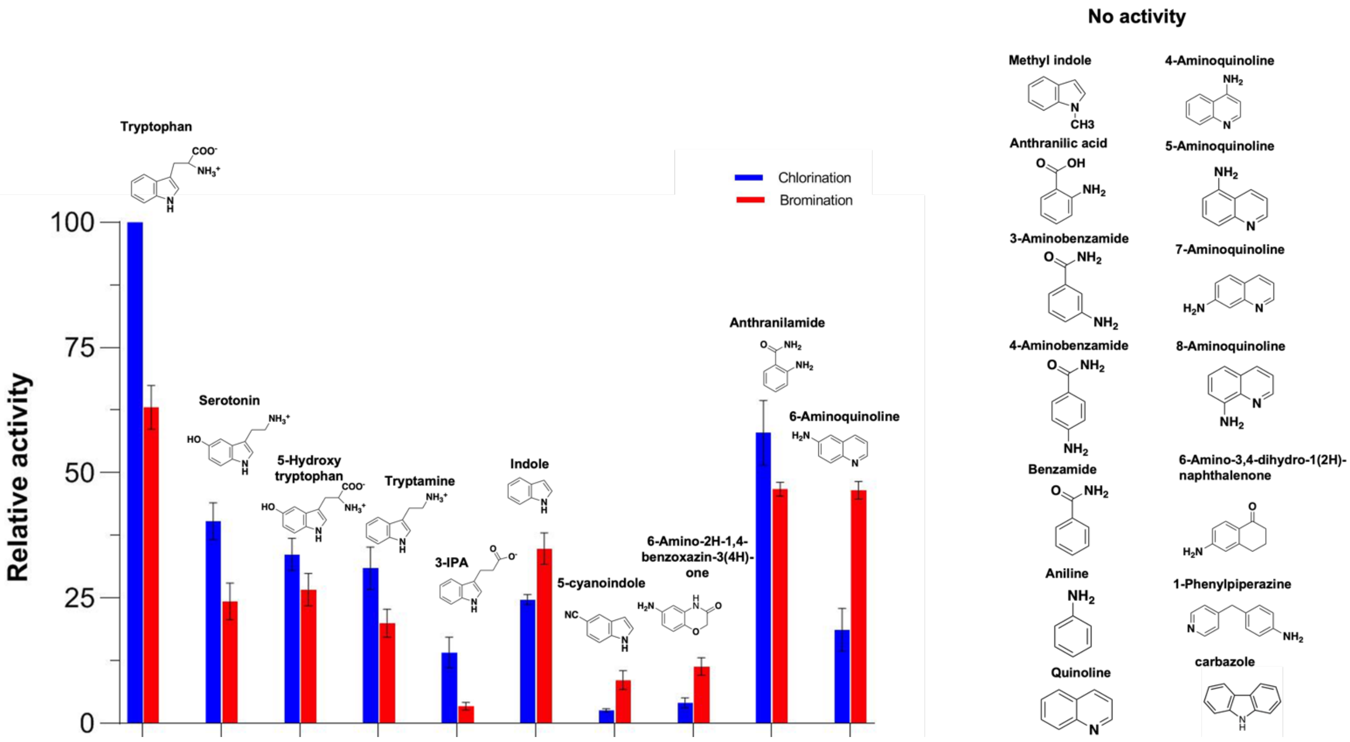


**Figure S6. BorH chlorinates and brominates non-tryptophan aromatic substrates.**

Results from *in vitro* chlorination and bromination assays of AbeH with Trp and 20 other aromatic substrates under identical reaction conditions.

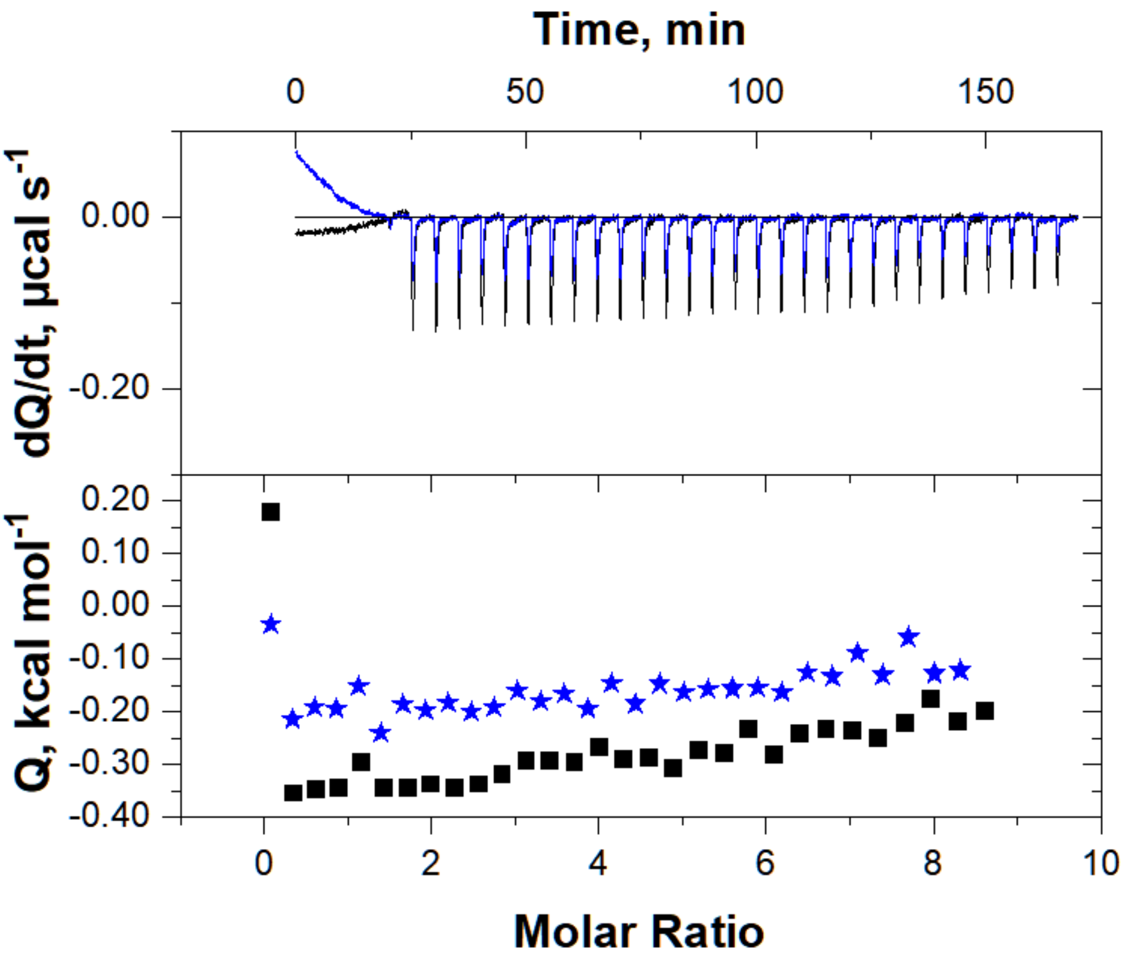


**Figure S7: No binding detected by ITC when Trp is titrated into AbeH at low molar ratios.**

Top graph is the raw differential thermogram of titration with baseline as a solid line, and the bottom graph is the integrated heat for each injection. Trp → buffer (heat of dilution; blue) and Trp → AbeH (black) are both plotted. There is no significant heat of binding detected at Trp:AbeH molar ratios between 0-9. Compare to Figure 5A, D.

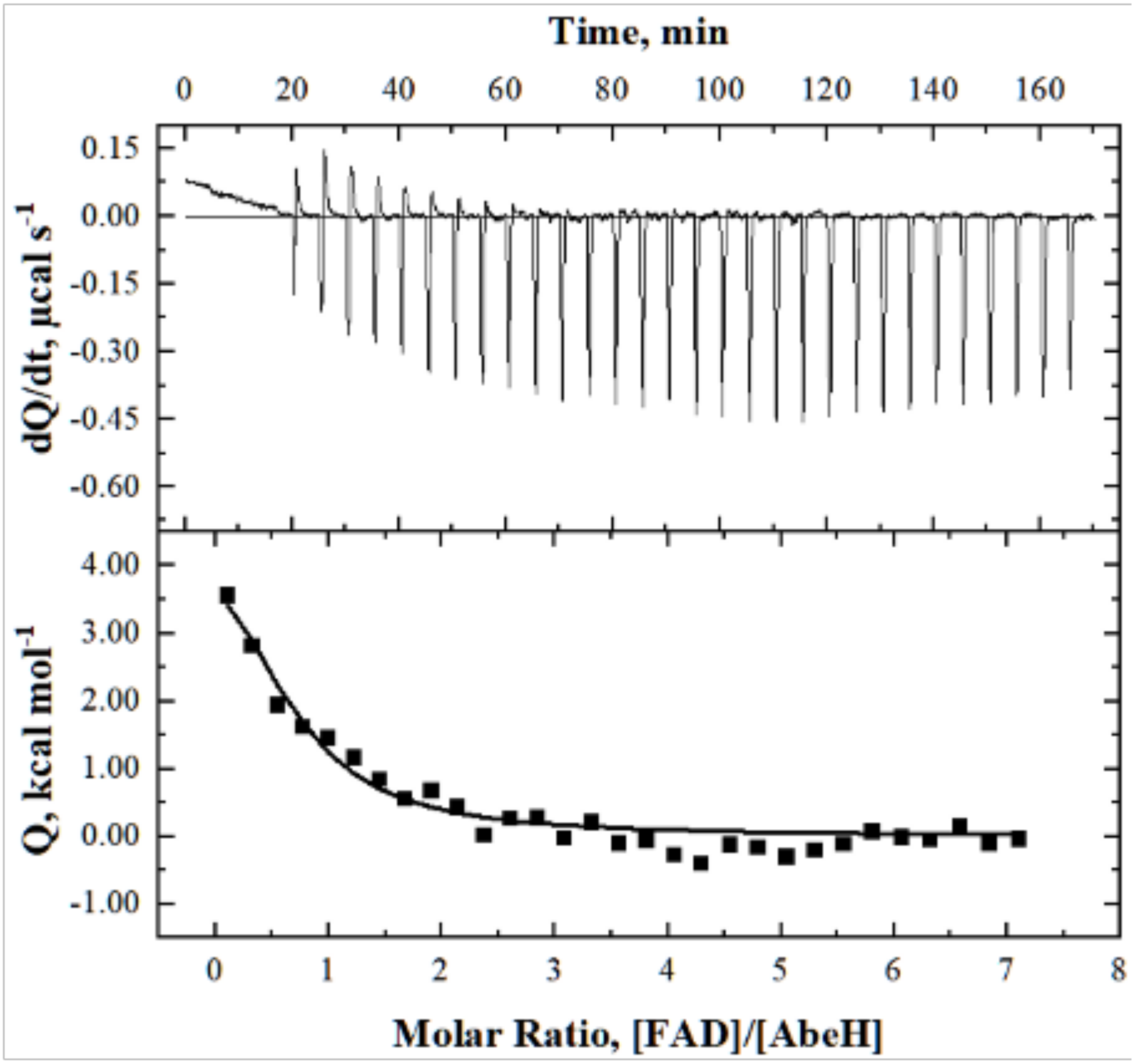


**Figure S8: Confirmation of endothermic titration of FAD into AbeH/Trp.**
Top graph is the raw differential thermogram of titration with baseline as a solid line, and the bottom graph is the integrated titration curve showing corrected heat for each injection (▪) and best fit line (–). This second experiment confirmed the data in Fig. 4C, showing an endothermic binding event. This experiment was carried out with a 2x higher molar ratio of Trp:AbeH than the experiment showing in Figure 4C, and the $K_D$ was higher (6.3 µM compared to 2.1 in Figure 4C).

**A**. BorH/FAD **B**. AbeH/FAD

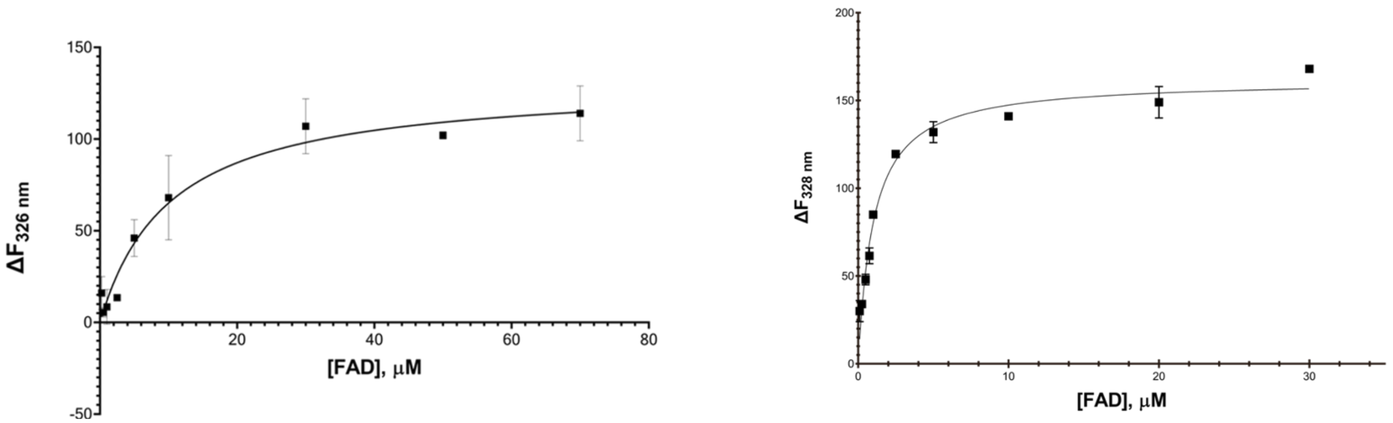


**Figure S9: FAD binding determined by quenching of BorH and AbeH intrinsic Trp fluorescence.**

**A**. Variable concentrations of FAD (0-70 μM) were incubated with 100 nM BorH and Trp fluorescence was measured with excitation at 290 nm and emission at 326 nm. Experiments were carried out in duplicate and mean values were fit to equation 1 using non-linear regression with GraphPad Prism 8.0, determining $K_D$ for BorH/FAD = 9.1 μM

**B**. Variable concentrations of FAD (0-30 μM) were incubated with 100 nM AbeH and Trp fluorescence was measured with excitation at 290 nm and emission at 326 nm with excitation at 295 nm and emission at 328 nm. $K_D$ for AbeH/FAD = 0.96 μM.

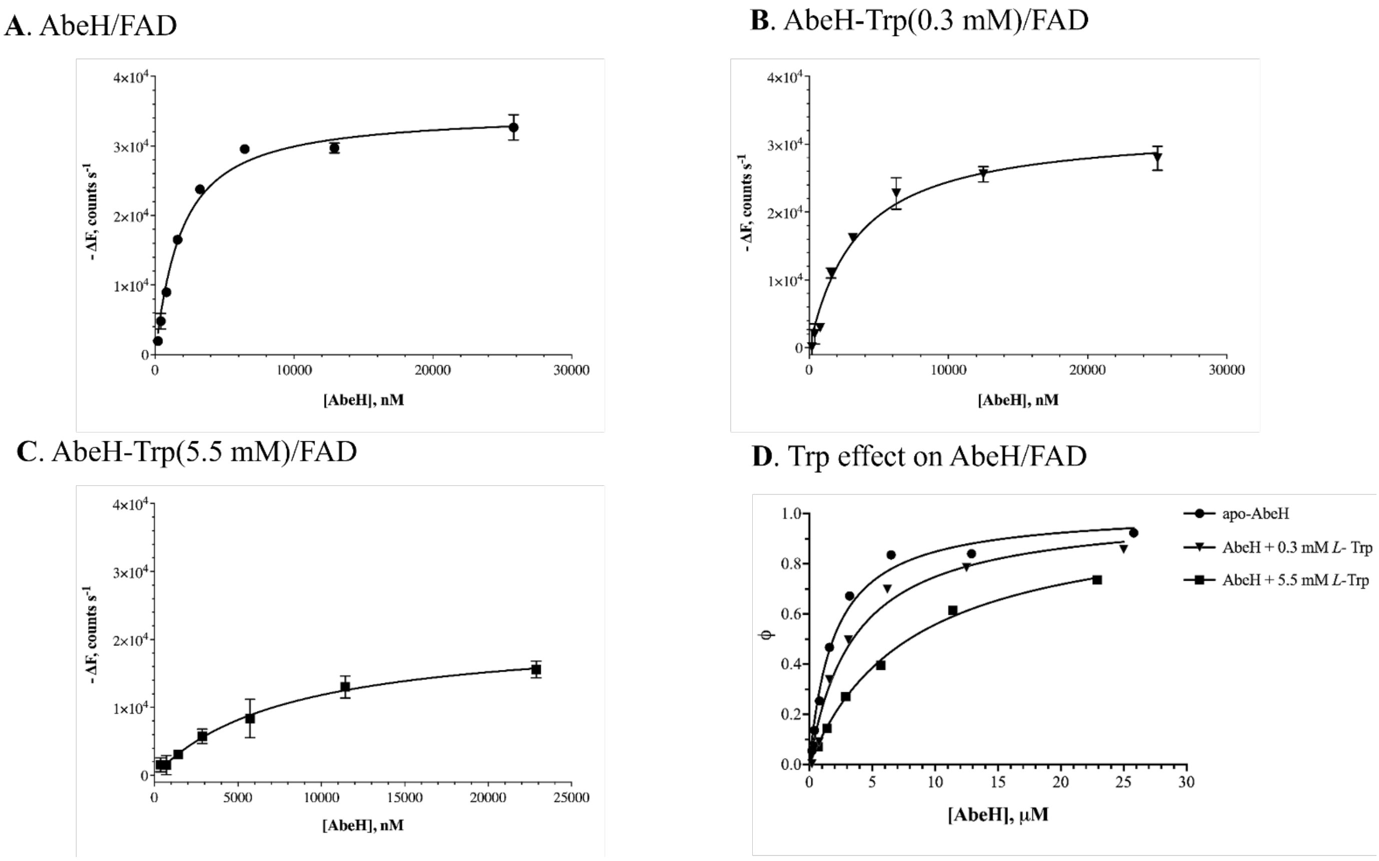


**Figure S10: Trp increases the $K_D$ for FAD binding to AbeH, providing cross-validation of negative cooperativity.**

Variable concentrations of AbeH or AbeH preincubated with Trp were added to and incubated with constant FAD and FAD fluorescence was measured with excitation at 450 nm and emission at 525 nm. Experiments were carried out in duplicate and mean values were fit to equation 1 using non-linear regression with GraphPad Prism 8.0.

A. Fluorescence quenching observed from binding of Apo-AbeH (0.20-26 μM) to 0.50 μM FAD; AbeH/FAD $K_D$ =1.6 μM in the absence of Trp.

B. Fluorescence quenching observed from binding of AbeH (0.20-25 μM) preincubated with 0.30 mM Trp to 0.49 μM FAD; AbeH/FAD $K_D$ = 3.1 μM in the presence of 0.3 mM Trp

C. Fluorescence quenching observed from binding of AbeH (0.40-23 μM) preincubated with 5.5 mM Trp to 0.45 μM FAD; AbeH/FAD $K_D$ = 7.7 μM in the presence of 5.5 mM Trp.

D. Fractional saturation (Φ) was calculated from equation (1) by the relation of $\Delta F/\Delta F_{max}$ and plotted against AbeH concentration to superimpose results of A-C.

**Table S2: ITC determined thermodynamic parameters of FAD, $FADH_2$, and Trp binding to AbeH and BorH.** Values were obtained from non-linear least squares fit analysis of the ITC data using the one-site model in Origin 7.0.

| **ITC Syringe** | **ITC Cell** | ***n*** | $K_A$ ($M^{-1}$) | $K_D$ ($\mu M$) | ***c*** | $\Delta H$ (kcal $mol^{-1}$) | $-T\Delta S$(kcal $mol^{-1}$) | $\Delta G$ (kcal $mol^{-1}$) | $\chi^2/DoF$ |
|---|---|---|---|---|---|---|---|---|---|
| Trp | AbeH | Fixed at 1 | $(1.2 \pm 0.2) \times 10^3$ | 833 | 0.02 | -4.7± 0.4 | 0.4 | -4.3 | 39 |
| Trp | BorH | Fixed at 1 | $(2.3 \pm 0.3) \times 10^3$ | 435 | 0.02 | -4.1± 0.3 | -0.5 | -4.6 | 8.8 |
| FAD | AbeH | 0.93 ± 0.05 | $(8.5 \pm 3) \times 10^5$ | 1.2 | 14 | -4.6 ± 0.4 | -3.6 | -8.2 | $5.2 \times 10^4$ |
| FAD | BorH | 1.1 ± 0.4 | $(7.8 \pm 1.9) \times 10^4$ | 13 | 0.7 | -11.4 ± 0.4 | 4.7 | -6.7 | $3.8 \times 10^4$ |
| $FADH_2$ | AbeH | 0.82 ± 0.03 | $(8.7 \pm 4.5) \times 10^6$ | 0.12 | 78 | -15.3 ± 0.7 | 5.7 | -9.6 | $1.1 \times 10^6$ |
| $FADH_2$ | BorH | 0.97 ± 0.05 | $(4.9 \pm 1.8) \times 10^6$ | 0.21 | 14 | -24.2 ± 1.6 | 15 | -9.2 | $2.4 \times 10^6$ |
| Trp | AbeH/FAD | No binding | | | | | | | |
| Trp | BorH/FAD | No binding | | | | | | | |
| Trp | AbeH/$FADH_2$ | No binding | | | | | | | |
| Trp | BorH/$FADH_2$ | No binding | | | | | | | |
| FAD | AbeH/Trp (16 mM) | 0.58 ± 0.02 | $(4.7 \pm 0.9) \times 10^5$ | 2.1 | 7.6 | 7.3 ± 0.4 | -15.1 | -7.8 | $9.7 \times 10^4$ |
| FAD | AbeH/Trp (28 mM) | 0.6 ± 0.1 | $(1.6 \pm 0.6) \times 10^5$ | 6.3 | 1.5 | 5.9 ± 1.6 | -13.1 | -7.2 | $4.6 \times 10^4$ |
| FAD | BorH/Trp | No binding | | | | | | | |

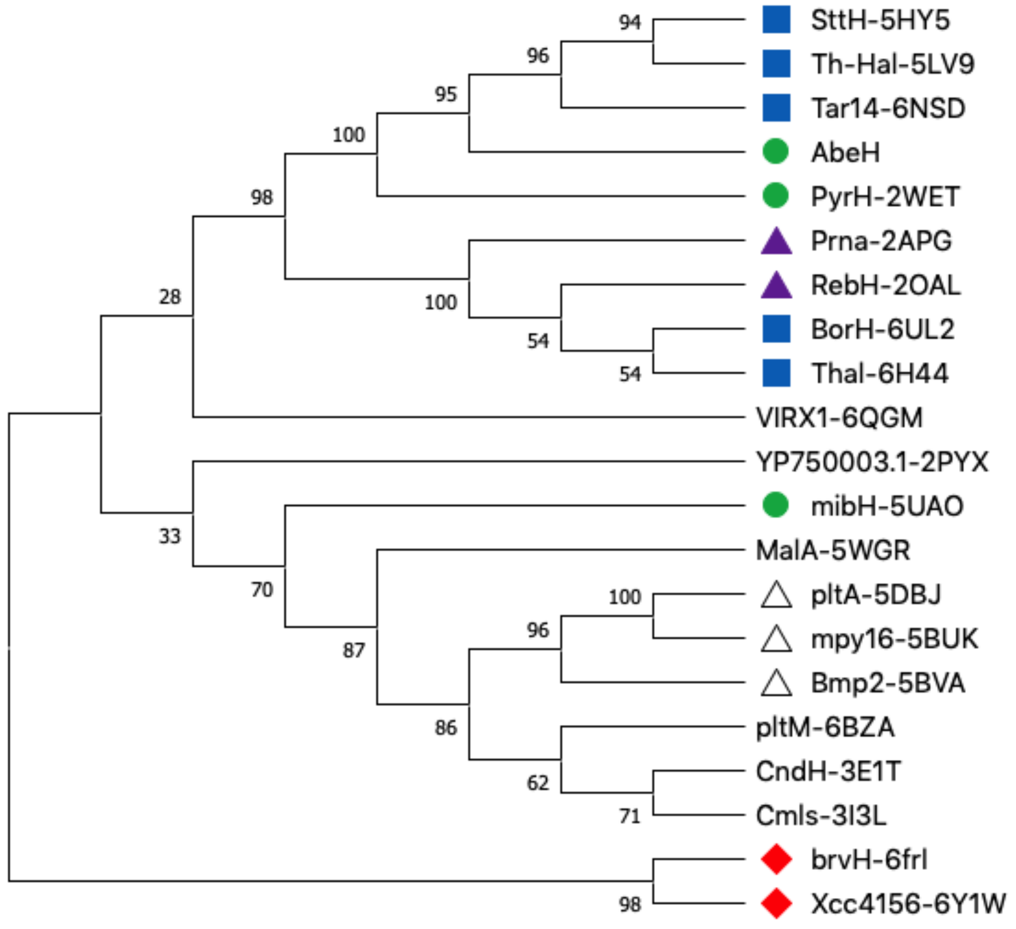


**Figure S11: AbeH and BorH belong to two different clades of FDHs**

Phylogenetic Bootstrap (100 run) consensus tree for evolutionary analysis by maximum likelihood method and JTT matrix-based model using neighbor-joining method with Mega X (1, 2). The percentage of this phylogenetic tree is shown next to branches, and associated taxa are clustered together. Trp-5-FDHs are labeled with green circles, Trp-6-FDHs are labeled with blue rectangles, Trp-7-FDHs are labeled with purple triangles, indole FDHs are labeled with red diamonds, and pyrrole halogenase are labeled with white triangles. PDB accession codes corresponding to the sequences used for the analysis are indicated.

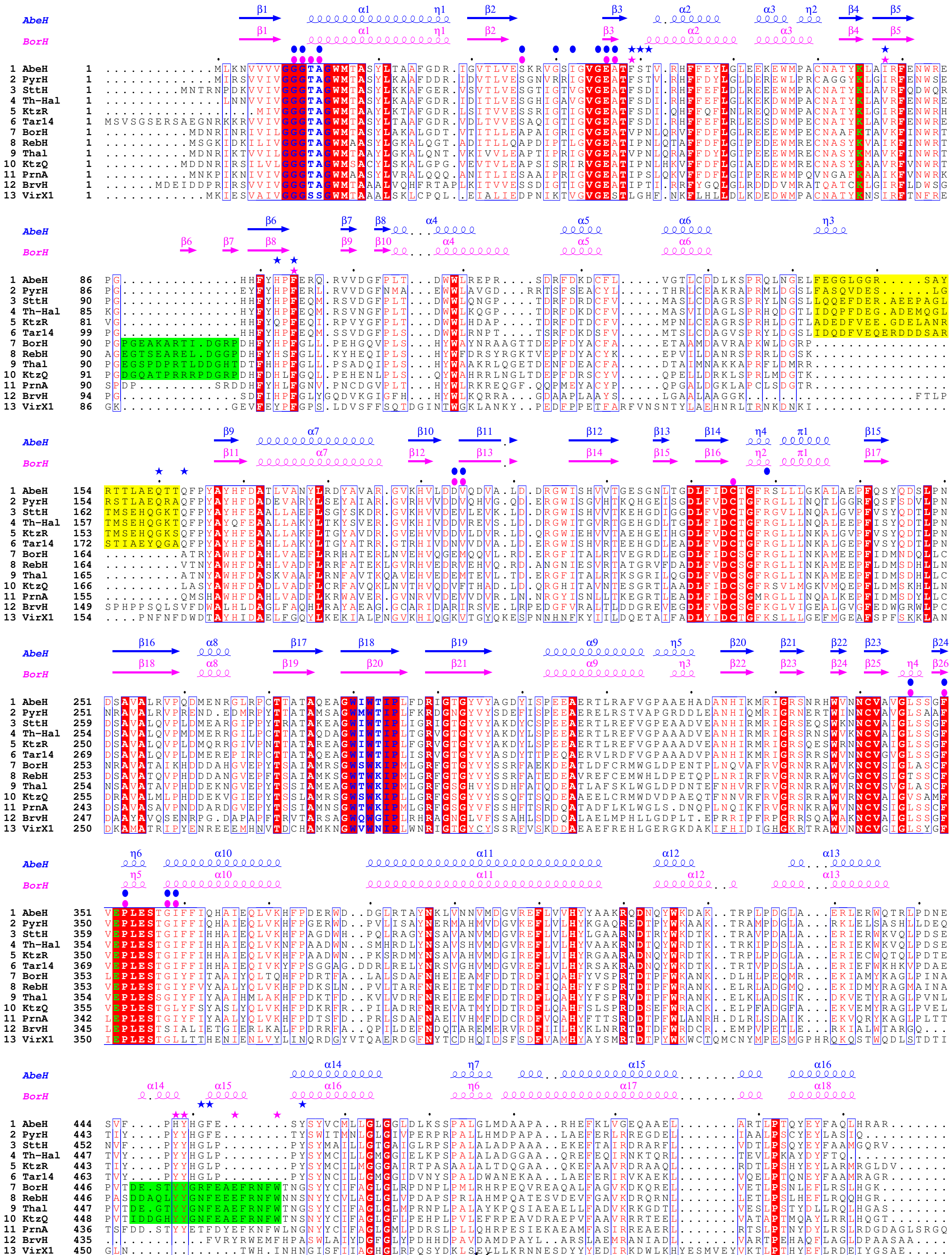
AbeH
BorH
1 AbeH
2 PyrH
3 SttH
4 Th-Hal
5 KtzR
6 Tar14
7 BorH
8 RebH
9 Thal
10 KtzQ
11 PrnA
12 BrvH
13 VirX1

**Figure S12: Multiple sequence alignment of AbeH and BorH with other FDHs.** Sequences were aligned with ClustalOmega(3) and alignment was formatted with ESPript(4). Secondary structure from AbeH/FAD/Cl- chain B (blue) and BorH/FAD chain A (pink) is displayed above the sequence alignment. Blue stars indicate residues contacting Trp in PyrH/Trp. Pink stars indicate residue contacting Trp in BorH/Trp. Blue ovals indicate residues contacting FAD in AbeH/FAD/Cl-. Pink ovals indicate residues contacting FAD in BorH/FAD/Cl-. The catalytic lysine (K75 in AbeH, K79 in BorH) and glutamate (E352 in AbeH, E354 in BorH) are in green. The insertion between $\alpha$6 and $\beta$9 unique to the AbeH clade is highlighted in yellow. The insertions unique to the BorH clade (between $\beta$5 and $\beta$6/8 and between $\alpha$13 and $\alpha$14/16) are highlighted in green. Residues in the conserved GxGxxG flavin-binding motif and the halogenase-specific WXWXIP motif are in blue. Strictly conserved residues are highlighted in red; highly conserved residues are boxed in blue.

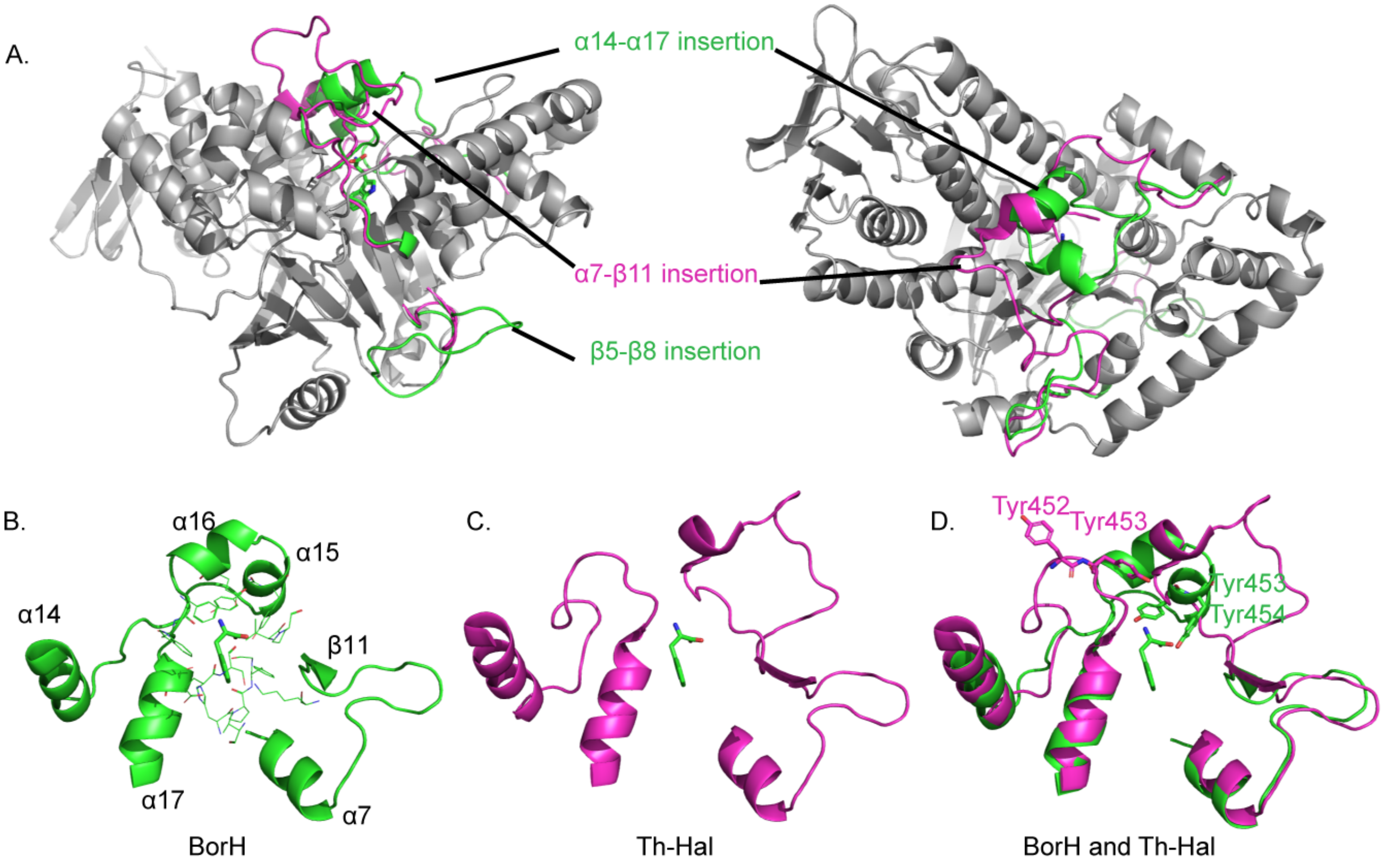


**Figure S13: Difference in Trp binding lid between BorH clade and AbeH clade (represented by Th-Hal.**

A. Ribbon diagram of BorH (gray) with three insertions highlighted. Green indicates the structure of BorH in these regions (which is the same in RebH and Thal), and magenta is the structure of Th-Hal (5LV9.pdb) in these regions (which is shared with AbeH, PyrH, SttH, and Tar14).

B. The Trp binding site lid in the BorH clade consists partly of residues from the α14-α17 insertion (including helices α15 and α16).

C. The Trp binding site lid in the AbeH clade lacks the α14-α17 insertion but has a long insertion between α7 and β11 that occupies the same space and interacts with the Trp in PyrH. Trp shown is from the BorH structure (The structure of Th-Hal does not have Trp bound).

D. Superposition of B and C, showing how the BorH clade insertion and the Th-Hal clade insertion generate a lid from topologically distinct structural elements. Interestingly, there is a YY sequence motif conserved in both the BorH clade and the AbeH clade in the region between α14 and α17 (Y453/454 in BorH and Y452/453) in Th- Hal), but these residues do not occupy the same positions in the structures.

**A.**

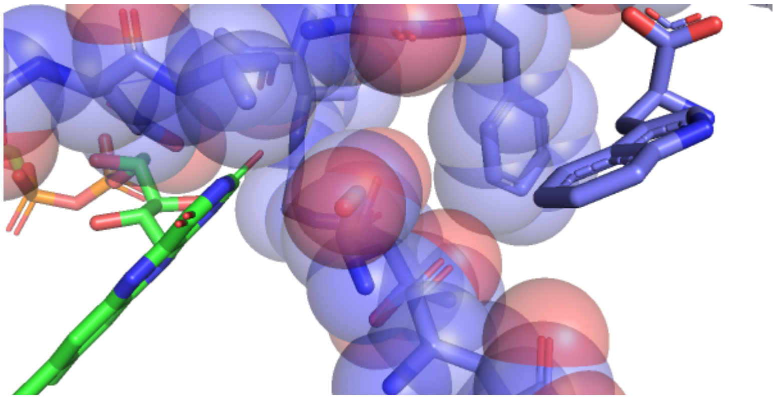

**B.**

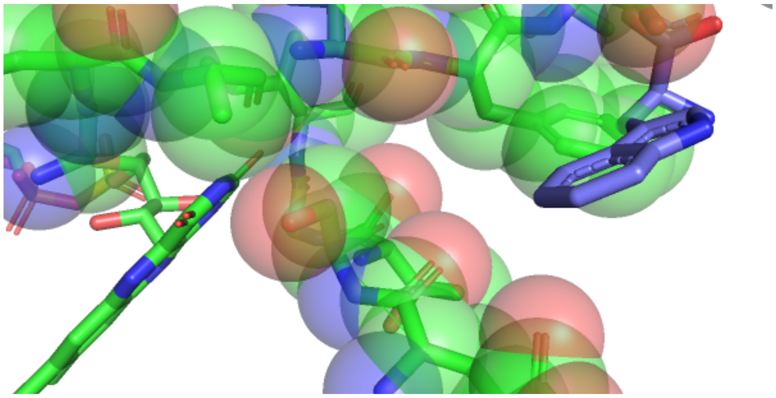

**Figure S14: Movement of the Trp gate upon FAD binding creates steric hindrance for Trp**

AbeH/FAD/$Cl^-$ (green carbon atoms) is superimposed with PyrH/Trp (purple carbon atoms)

A. PyrH/Trp with FAD from AbeH/FAD/$Cl^-$
B. With Trp from PyrH/Trp. Closing of flavin binding loop over FAD and flipping of the Glu46 switch pushes Ser49 and Phe50 into the Trp binding site, preventing Trp from binding due to steric collision.

A. B.

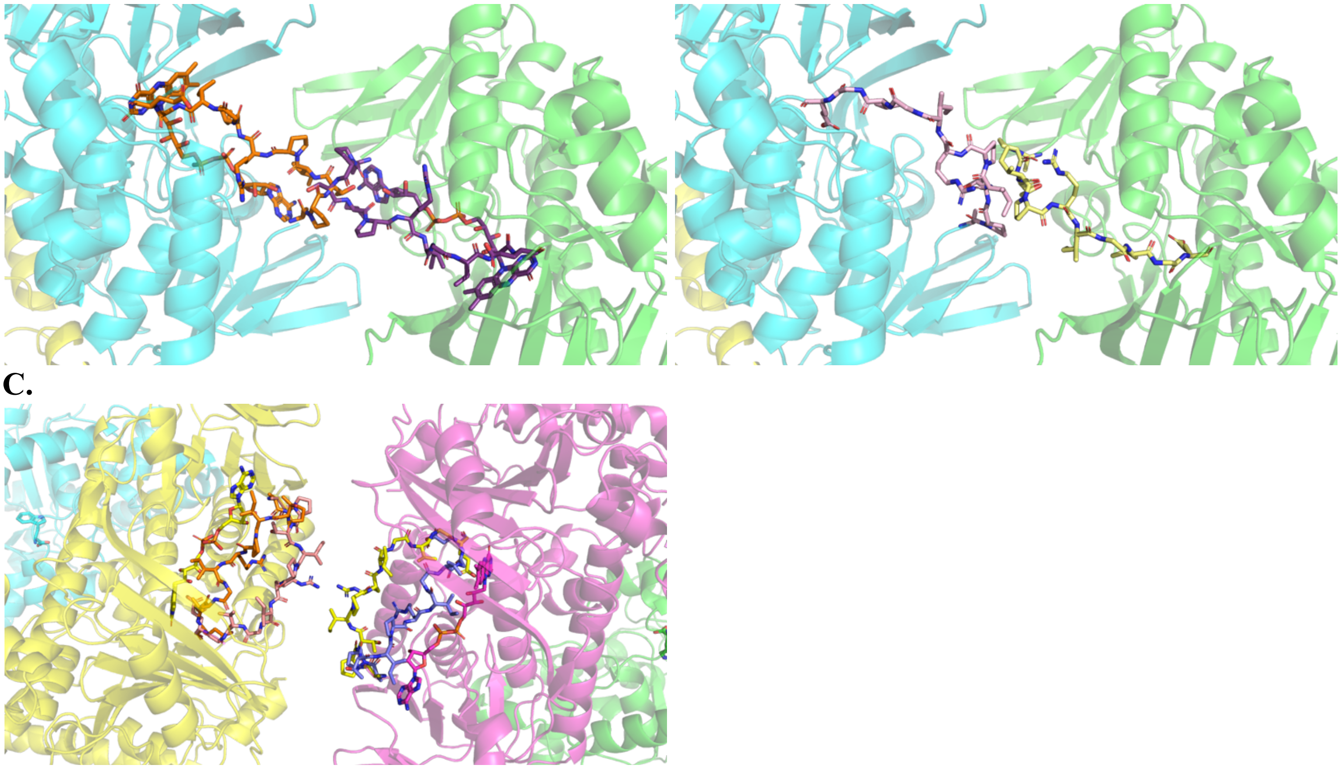

C.

Figure S15: Crystal contact preventing FAD binding in Chains A and B of BorH/Trp + BorH/FAD structures.

Steric clashes between BorH chains would results from either the open or closed conformation of the flavin binding loop. The proximity of the two chains at this contact and the steric hindrance for the functional flavin loop conformations explains why FAD could not be soaked into chains A and B, and why there is no interpretable density for the flavin binding loop.

A. Ribbon diagram of BorH/Trp Chain A (green) and BorH/Trp Chain B (cyan) interface between ASUs in the BorH/FAD+BorH/Trp crystal structure. Thal/FAD with the flavin binding loop in the closed conformation (7CU1A; purple and orange) has been superimposed over both chains of BorH and FAD and the flavin binding loops from those structures are shown. There are steric clashes between the loops and the adenosine portions of FAD.
B. Ribbon diagram of BorH/Trp Chain A (green) and BorH/Trp Chain B (cyan) interface between ASUs in the BorH/FAD+BorH/Trp crystal structure. Thal/AMP with the flavin binding loop in the open conformation (7CU1B; pink and yellow) has been superimposed over both chains of BorH and the flavin binding loops from those structures are shown.
C. Ribbon diagram of the interface between BorH/FAD Chain C (yellow) and BorH/FAD Chain D (magenta) in the BorH/FAD+BorH/Trp crystal structure. Thal structures with the flavin binding loop in the closed conformation (7CU1A Thal/FAD; purple and orange) and the open conformation (7CU1B Thal/AMP; pink and yellow) are superimposed over both chains of BorH. Both conformations of the loop can be accommodated without steric clashes.

**Table S3. Conformations of the flavin binding loop and Trp binding site observed in crystal structures of FDHs in the AbeH clade.**

| | Flavin-binding loop | "Switch" Glu | "Gate" F49/S50 | "Lid" 152-160 | "Lid" 161-167 | Sequence identity to AbeH (%) |
|---|---|---|---|---|---|---|
| **AbeH/FAD/ $Cl^-$ B** | Closed | Out | Closed | Disordered | Open | 100<br>100 |
| **Apo-AbeH C** | Open | In | Closed | Disordered | Open | |
| **PyrH/Trp (2WEUC)** | Open (disordered) | In | Open | Ordered | Closed | 62 |
| **PyrH/FAD/ $Cl^-$ (2WETC)** | Closed | Out | Closed | Disordered | Open | |
| **PyrH/FAD/ Trp/ $Cl^-$ (2WETB)** | Closed | Out | Open | Ordered | Closed | |
| **SttH/FAD/ $Cl^-$ (5HY5)** | Closed | Out | Closed | Disordered | Open | 68 |
| **Apo-Th-Hal (5LV9)** | Open | In | Open | Ordered | Closed | 67 |
| **Tar14/FAD (6NSD)** | Closed | Out | Closed | Ordered | Closed | 68 |

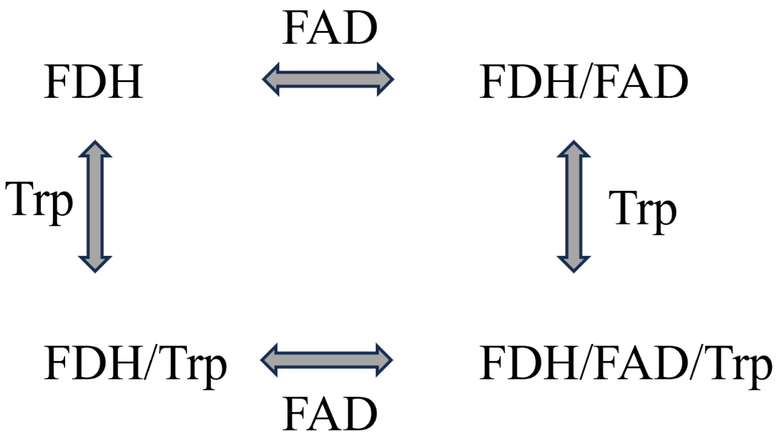


**Figure S16: ITC experiments designed to test ternary complex formation in FDH.** Both FDHs, AbeH and BorH were tested using this model described. This ITC based cooperativity model was elaborately discussed in the referenced literatures(5, 6).

Supporting Information References:

1. Kumar, S., Stecher, G., Li, M., Knyaz, C., andTamura, K. (2018) MEGA X: Molecular Evolutionary Genetics Analysis across Computing Platforms *Mol Biol Evol* **35**, 1547-1549 10.1093/molbev/msy096
2. Saitou, N., andNei, M. (1987) The neighbor-joining method: a new method for reconstructing phylogenetic trees *Mol Biol Evol* **4**, 406-425 10.1093/oxfordjournals.molbev.a040454
3. Sievers, F., andHiggins, D. G. (2014) Clustal Omega *Curr Protoc Bioinformatics* **48**, 3.13.1-3.13.16 https://doi.org/10.1002/0471250953.bi0313s48
4. Gouet, P., Courcelle, E., Stuart, D. I., andMétoz, F. (1999) ESPript: analysis of multiple sequence alignments in PostScript *Bioinformatics* **15**, 305-8 10.1093/bioinformatics/15.4.305
5. Brown, A. (2009) Analysis of cooperativity by isothermal titration calorimetry I*nt J Mol Sci* **10**, 3457-77 10.3390/ijms10083457
6. Velazquez-Campoy, A., Goñi, G., Peregrina, J. R., andMedina, M. (2006) Exact analysis of heterotropic interactions in proteins: Characterization of cooperative ligand binding by isothermal titration calorimetry *Biophys J* **91**, 1887-904 10.1529/biophysj.106.086561
